# Global Dynamics as Communication Sensors in Peptide Synthetase Cyclization Domains

**DOI:** 10.1101/2021.10.06.461881

**Authors:** Subrata H. Mishra, Aswani K. Kancherla, Kenneth A. Marincin, Guillaume Bouvignies, Santrupti Nerli, Nikolaos Sgourakis, Daniel P. Dowling, Dominique P. Frueh

**Author notes:** These authors contributed equally to this work (Aswani K. Kancherla, Kenneth A. Marincin). Reference Standard Laboratory, United States Pharmacopeial Convention, 12601 Twinbrook Pkwy, Rockville, MD, 20852, USA.

## Abstract

Structural biology is the foundation for deriving molecular mechanisms, where snapshots of macromolecules and binding partners inform on mutations that test or modify function. However, frequently, the impact of mutations violates the underpinnings of structural models, and mechanisms become cryptic. This conundrum applies to multidomain enzymatic systems called nonribosomal peptide synthetases (NRPSs), which assemble simple substrates into complex metabolites often with pharmaceutical properties. Engineering NRPSs can generate new pharmaceuticals^1-3^ but a dynamic domain organization challenges rational design.^4-8^ Using nuclear magnetic resonance (NMR), we determined the solution structure of a 52 kDa cyclization domain and demonstrate that global intra-domain dynamics enable sensing of substrates tethered to partner domains and draw an allosteric response encompassing the enzyme’s buried active site and two binding sites 40 Å apart. We show that a point-site mutation that impedes the domain dynamics globally hampers the allosteric response. We demonstrate this mechanism through NMR experiments that provide atomic-level read-outs of allosteric responses during biochemical transformations *in situ*. Our results establish global structural dynamics as sensors of molecular events that can remodel domain interactions and illustrate the need for integrating structural dynamics explicitly when deriving molecular mechanisms through mutagenesis and structural biology.

## INTRODUCTION

Structural dynamics are increasingly recognized as a component of biological activity yet determining their presence and their function remains challenging. Structural dynamics play a role in enzymatic activity,^9-12^ allosteric communication,^13-15^ and molecular binding.^16^ However, detecting dynamics and determining their function is experimentally demanding, often because establishing correlations between complementary experiments is hindered by experimental precision. Notably, although NMR can provide valuable information, the method is challenged by molecular size. Yet, when dynamics go undetected, kinetic models are incomplete, allostery may go unnoticed, and structure-based mechanisms often need revisions. The latter is true for large enzymatic systems called nonribosomal peptide synthetases, and we sought to determine whether structural dynamics may resolve gaps in understanding these fascinating enzymes.

NRPSs are microbial molecular factories that employ a dynamic multi-domain architecture to assemble simple substrates into complex natural products, including pharmaceuticals such as antibiotics (bacitracin), anticancer agents (epothilones), or immunosuppressants (cyclosporins).^17^ Adenylation (A) domains attach substrates to 20 Å phosphopantetheine moieties (Fig. 1a) of holo thiolation (T) domains through thioester bonds (Fig. 1b). Condensation (C) domains then catalyse the peptide bond formation between substrates of upstream and downstream T-domains, leading to a downstream acceptor harbouring an extended intermediate and an upstream donor restored to its holo form (Fig. 1c). As part of the C-domain family, cyclization (Cy) domains further catalyse cyclodehydration (Fig. 1d). Iteration leads to chain elongation before product release, e.g. through thioesterases. Our model system, HMWP2, combines salicylate with two cysteines to synthesize a precursor of yersiniabactin (Fig. 1e), a virulence factor of *Yersinina pestis*, the causative agent of the bubonic plague. Such modular organization has stimulated efforts to engineer NRPSs and control substrate incorporation to produce novel pharmaceuticals^1-3^, but a dynamic domain architecture^4-8^ hampers rational design. Notably, T-domains visit catalytic domains through sequential, transient interactions (Fig. 1f), and it is unclear whether these interactions are random or if chemistry promotes interactions in line with synthesis. Further, intra-domain structural dynamics are often necessary for function,^18,19^ although dynamics within condensation domains have proven puzzling. Schmeing and Co. noted that open and closed conformations observed in C-domains may be needed at different stages of synthesis,^20^ but the conformations of C-domains in crystal structures are unaltered in the presence of other domains or substrates (Supporting Information (SI) Fig. 1). The mechanisms of peptide bond formation and heterocyclization are under frequent revisions and mutagenesis impacts function in sometimes unexpected manner.^21^ Thus, the C-domain family appears to bear the hallmarks of structural dynamics, and we strived to determine them experimentally and establish their function using the domain Cy1 of HMWP2.

**Figure 1.**
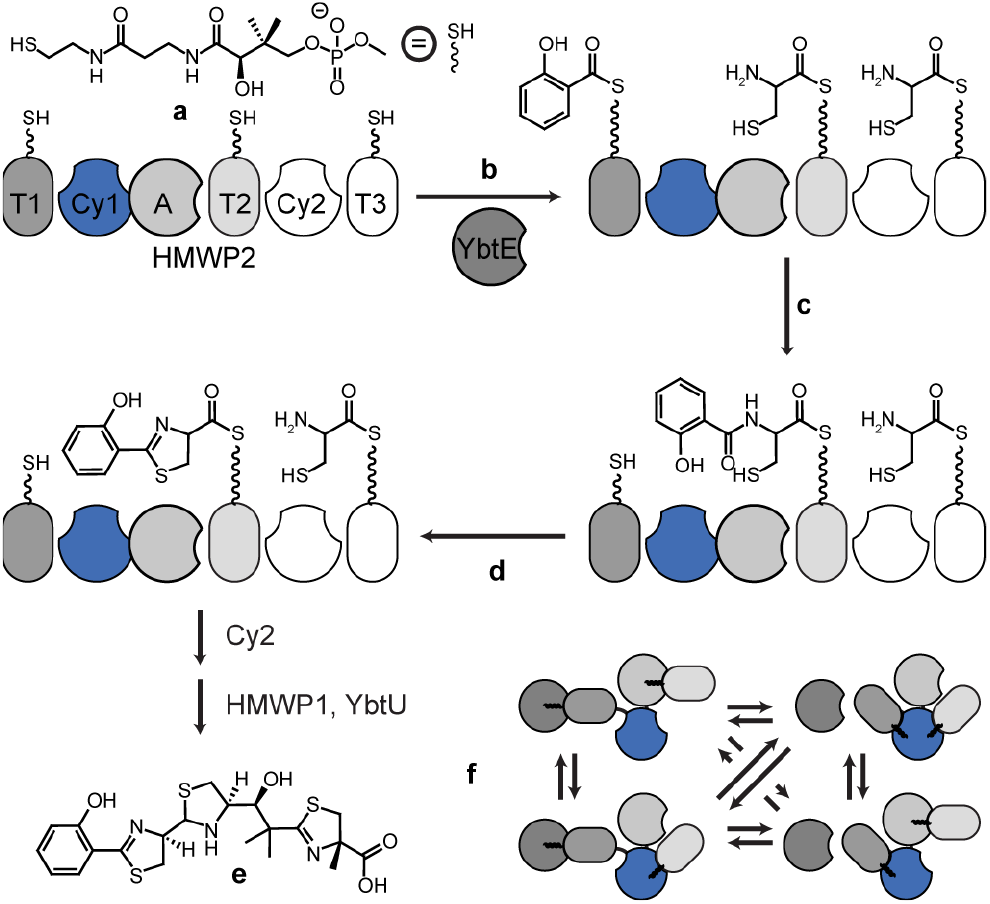
HMWP2 and NRPS synthesis. **a**, holo T-domains harbor phosphopantetheines. **b**, Adenylation domains (*in cis*, A, and *in trans*, YbtE) load substrates onto T-domains. **c**, C- (not shown) and Cy-domains catalyze peptide bond formation between substrates of T domains. **d**, Cy-domains additionally catalyze cyclodehydration. **e**, Cy2, HMWP1, and YbtU finalize the synthesis of yersiniabactin. **f**, Selected domain interactions during NRPS synthesis involving T, A and Cy/C domains.

## RESULTS AND DISCUSSION

### Structural fluctuations in Cy1

The solution structure of Cy1 displays the conserved C-domain family fold^21^, but its NMR ensemble reflects malleability (SI Table 1, Fig. 2, SI Fig. 2). N- and C-terminal residues display distinct connected regions, with C-terminal residues (Fig. 2a, dark blue) spatially crossing over twice into the N-terminal region (Fig. 2a, c). Both crossovers include loops (L16 and L20, SI Fig. 3) that interact with the donor T-domain. L20 leads to an extended region that occasionally pairs with β-strand β2 of the N-terminal region in the NMR ensemble. Like all Cy- and C-domains, Cy1 displays a tunnel between both regions, which harbours the active sites for peptide bond formation^21^ and cyclodehydration.^22,23^ In multi-domain NRPS structures, donor and acceptor T-domains funnel their phosphopantetheine arms from opposite ends of this 40 Å tunnel (Fig. 2a, pink, purple) to reach the buried active site.^8,24,25^ NMR provides ensemble of conformers (SI Fig. 2) that may reflect dynamics. Most notably, Cy1’s tunnel is constricted in several conformers, suggestive of transient interactions within the tunnel (Fig. 2d, SI Fig. 2c). Further, the tunnel entrances at the acceptor (Fig. 2b) and donor (Fig. 2c) sites display structural heterogeneity that occasionally prevent substrate access. This malleability recalls that seen in a distant member of the C-domain family, in which accelerated molecular dynamics revealed the closing of an otherwise open channel replacing the tunnel of this domains.^26^ Our results may reflect structural fluctuations, in which access to the tunnel is intermittent, a gating mechanism that can be harnessed for substrate recognition and that we investigated further.

**Figure 2.**
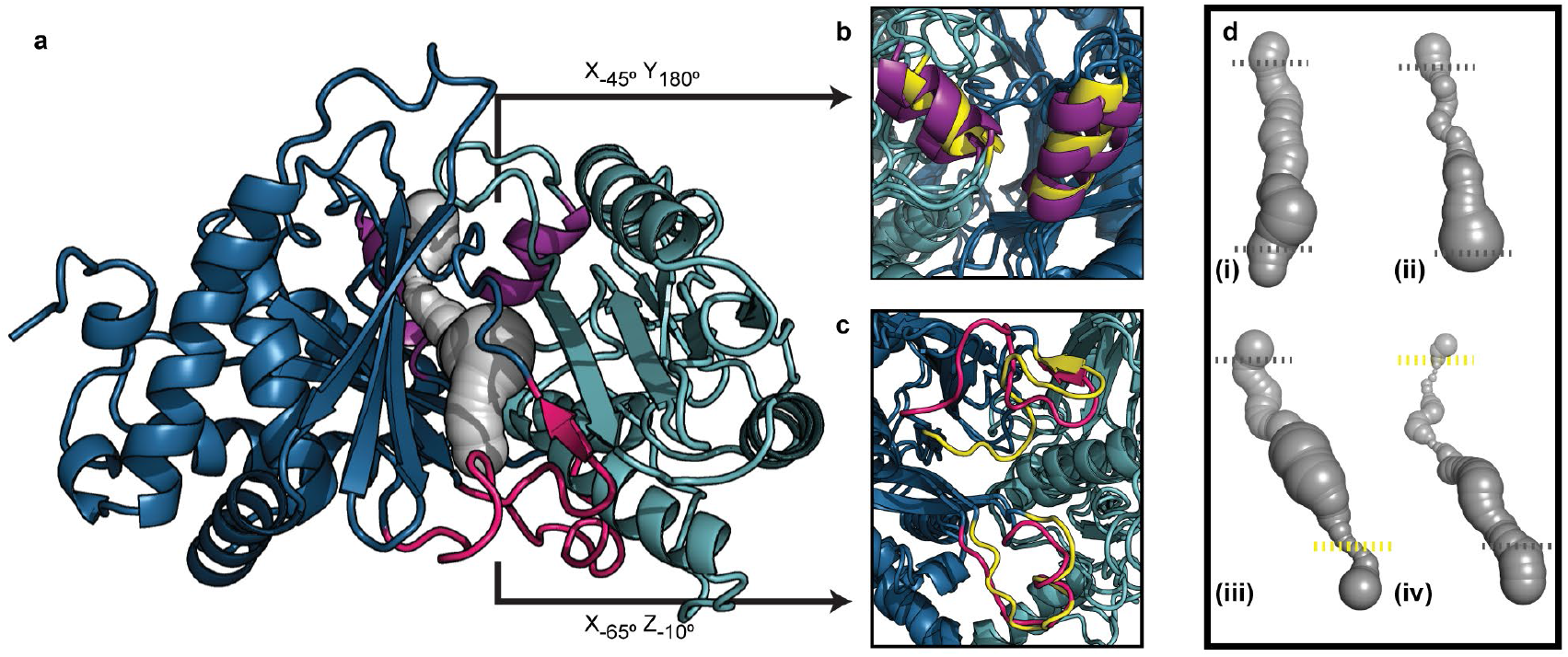
Solution structure of Cy1 and transient tunnel accessibility. **a**, medoid NMR conformer displayed with N- and C-subdomains in light and dark blue respectively. T domain sites in pink (donor) and purple (acceptor). **b, c**, open (purple, pink) and closed (yellow) conformations at acceptor (b) and donor (c) sites. **d**, tunnels in conformers are found fully open (i), constricted at the center (ii), closed at donor (iii), and closed at acceptor (iv) sites. Dashes approximate tunnel entrances (grey: open; yellow: constricted).

Cy1 is subject to global structural fluctuations at μs-ms time-scales, which mirror conformational changes in the C-domain family. We used NMR relaxation dispersion^27^ to identify residues that sample different chemical environments (Fig. 3a-b). We found that both T-domain binding sites are dynamic (Fig. 3a), consistent with structural fluctuations seen in our NMR ensemble and in crystallographic studies (SI Fig. 4) thus strengthening a gating mechanism where structural fluctuations modulate entry of the tethered substrates to access the active site. However, we also observed wide-spread dynamics involving 131 residues, with the N-terminal lobe displaying a dynamic footprint reminiscent of conformational changes reported by crystallography (Fig. 3b, c).^20^ Hence, we used multi-field relaxation dispersion to determine whether motions in Cy1 reflected global changes seen in C domains. The majority of dynamic residues (126/131) fit a global two-conformer model, with a minor conformer populated at 3% and an exchange rate of 1,522 s^-1^. Chemical shift analysis demonstrates that we are not probing unfolding of Cy1 (SI Fig. 4e, f). The function of this malleability appears to include a gating mechanism for substrate recognition, but the global nature of Cy1 dynamics also points at allostery, with a network of dynamic residues linking the remote T-domain binding sites. Thus, Cy1 dynamics may serve as a conduit to propagate the impact of T-domain binding from one site to the other remote site. We proceeded to demonstrate this hypothesis experimentally.

**Figure 3.**
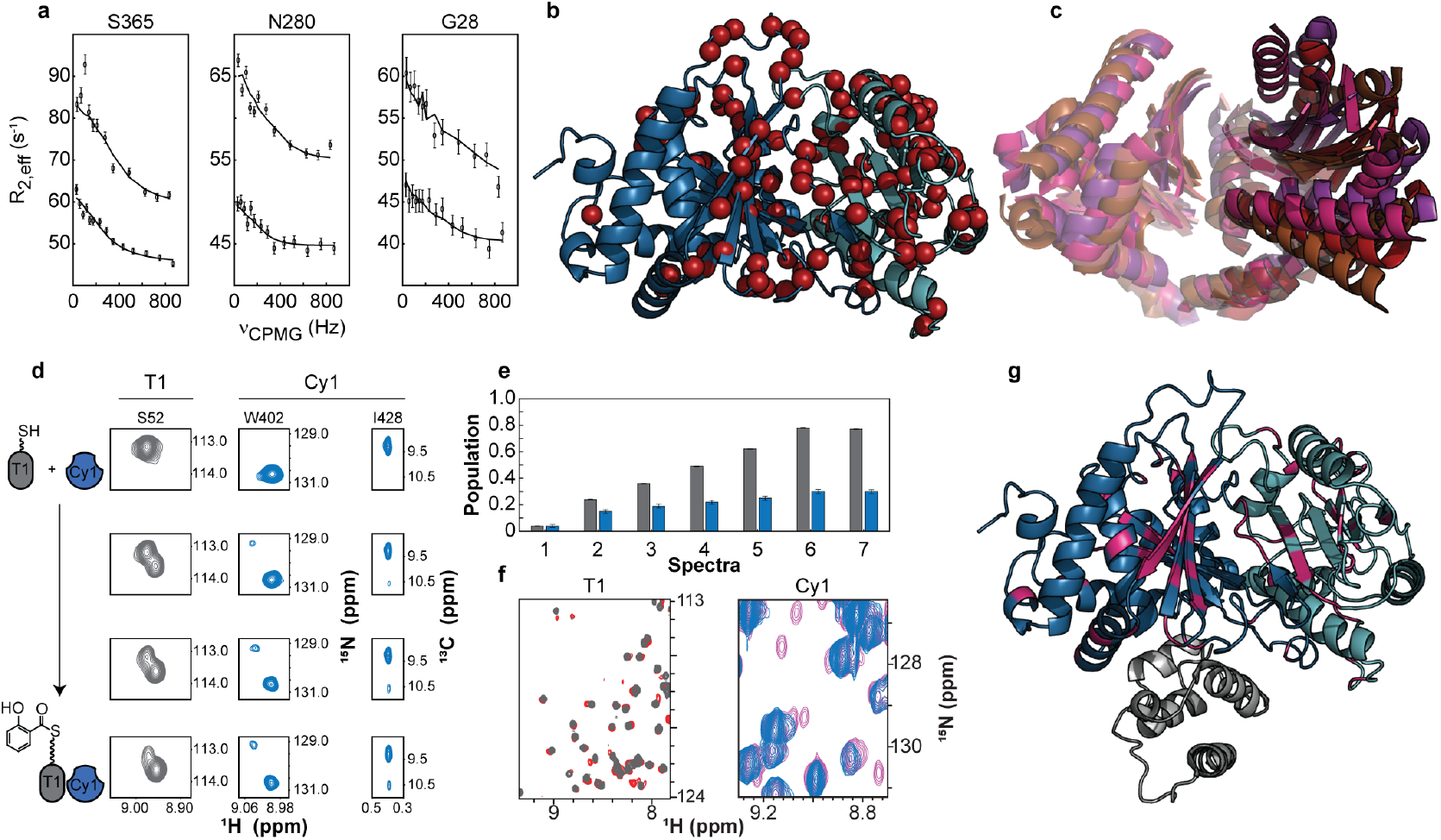
Dynamics and allosteric response in Cy1. **a**, Representative relaxation dispersion profiles of residues at the donor site (S365), within the tunnel (N280), and at the acceptor site (G28). **b**, Cy1 residues (red spheres) exhibiting dynamics on the ms-μs timescale. **c**, Crystal structures of C and Cy domains aligned by the C-terminal subdomain. (PDB: 5T3D, 4JN3, 6P1J, and 1L5A). For clarity, loops have been hidden. **d**, Time- shared NMR data showing concomitant loading of T1 (S52, grey) and emergence of a minor Cy1 population (W402 amide, I428 methyl, blue). **e**, Population of loading for T1 (grey) and of Cy1 minor conformation (blue). **f**, IDIS-NMR of T1 (holo:grey, loaded: red) and Cy1 spectra (free: blue, with T1 loaded: pink) allow for complete analysis of each protein without spectral crowding. **g**, Cy1 residues (pink) exhibiting a response to substrate-loaded T1.

### Cy1 responds allosterically to its partner substrate loading

We found that Cy1 responds to its donor thiolation domain only when it holds a substrate. Structures in the C-domain family appear unaffected by partner T-domains or substrates (SI Fig. 1), and substrates rapidly fall off of T domains, hampering traditional structure-function approaches to investigate allostery. We designed an experiment that builds upon our previous method^28^ to overcome these obstacles. We first presented Cy1 with its donor T-domain (T1) in holo form and observed no spectroscopic perturbation (SI Fig. 6a), suggestive of a dissociation constant larger than several mM. We then loaded T1 with its salicylate substrate *in situ* and used a combination of isotope labelling and tailored pulse sequences^29,30^ to monitor the spectra of each protein simultaneously. New Cy1 signals appeared as T1 was loaded with salicylate, denoting strong interaction between Cy1 and loaded T1 (Fig. 3d-f, SI Fig. 5). These observations corroborate recent studies where substrate-loaded donor T domains were needed to detect binding with a C-domain.^31^ Our results are remarkable as one would expect substrate recognition to take place at the active site near the centre of the tunnel, but our data indicates that the phosphopantetheine arm of holo-T1 does not go into the tunnel for substrate probing. Instead, our results point at substrate recognition at the surface of Cy1 paired with a dynamic gating mechanism, where dynamics respond to the presence of substrate to promote engagement.

Substrate recognition at the donor site induces allosteric remodelling of Cy1’s active site, tunnel, and remote acceptor binding site. New signals observed in Cy1 identify residues with altered environments when Cy1 binds to T1. Mapping affected residues (Fig. 3f, Cy1) on our structure (Fig. 3g) reveals not only changes in the donor site but depicts a conformer with an average population of approximately 11% (SI Fig. 5c) with changes encompassing the active site and the remote acceptor site. This conformer appears not to be selected from conformations of free Cy1 (SI Fig. 6). These observations demonstrate that, upon substrate recognition at the donor site, Cy1 responds allosterically to propagate changes to all sites involved in synthesis. Such a global reaction is reminiscent of Cy1’s global dynamics, and we next used mutagenesis to demonstrate that dynamics mediate this allosteric response.

### Impeding dynamics hinders recognition

Mutating a residue at the centre of the tunnel impairs both Cy1 dynamics and its response towards its substrate loaded partner. To demonstrate that dynamics facilitate Cy1’s allosteric response, we mutated a residue distant from the donor site, such that the mutation could not impact binding through direct interactions. The conserved aspartate D391 was shown by two teams to govern heterocyclization through interactions with the acceptor substrate.^22,23^ Unexpectedly, in Cy1, mutating this residue to an asparagine (D391N) induces a spectacular global response that reaches both binding sites (Fig. 4a, SI Fig. 7a). Such changes could indicate assaults to structural integrity but instead this mutation stabilizes Cy1’s structure (Fig. 4b, SI Fig. 8a-c) and impairs structural fluctuations, as evidenced by Hartman-Hahn R_ex_ profiles^32^ (Fig. 4c, SI Fig. 8d). The global response imparted by this mutation does not reflect conformational selection of a minor Cy1 conformation (SI Figs. 7b, c). Finally, we repeated our experiment to monitor allostery and, although a higher population of loaded T1 should have favoured complex formation for D391N (SI Fig. 9c), we observed a severely hampered response of D391N to loaded T1, with only five signals detected for D391N versus 77 for wild type Cy1 and a dramatic reduction in the apparent population of molecules responding (Fig. 4d). D391N’s mitigated response to loaded T1 indicates that, while the two domains do still interact, the formation of a minor state with an allosteric response is now hindered. These experiments establish dynamics as a mediator of the substrate-specific allosteric response we uncovered in the cyclization domain.

**Figure 4.**
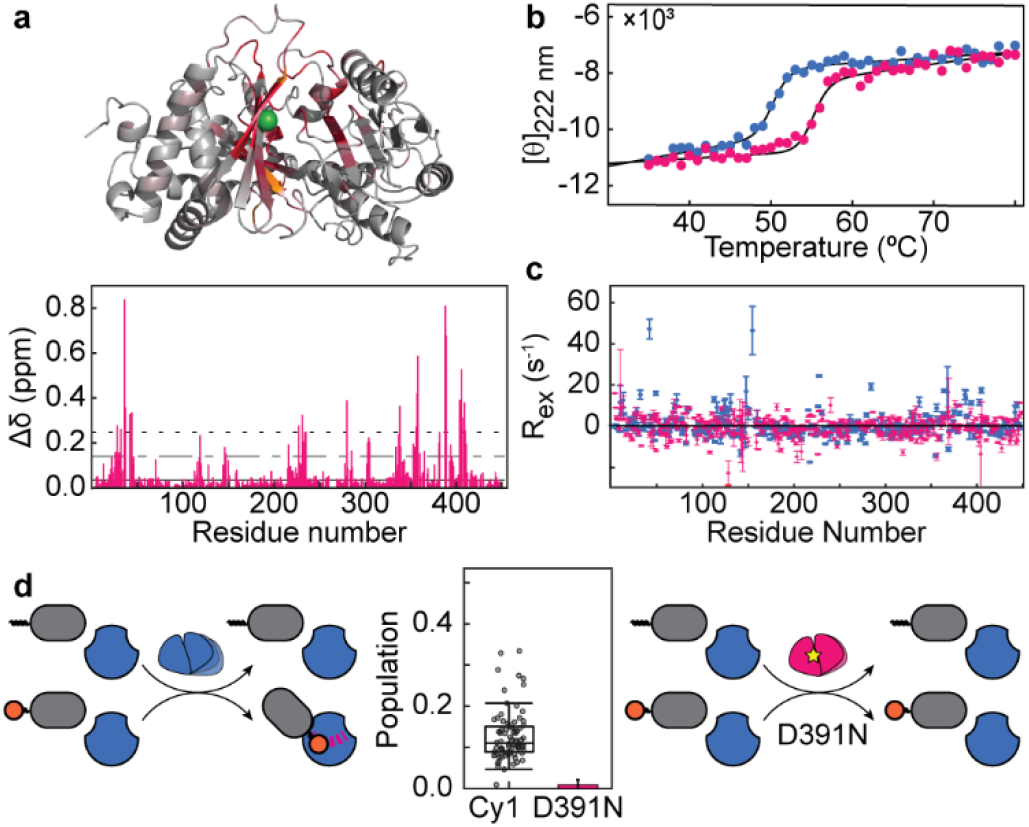
A Cy1 point-site mutation impedes dynamics and allosteric response. (a) Chemical shift perturbations (Δδ) between Cy1 and D391N mapped onto the Cy1 structure (top panel) using a color gradient where bright red indicates a CSP two standard deviations above the median (dotted line in the bottom panel). Residues that were uniquely assigned in D391N but not in wild type are in orange and D391 is shown as a green sphere (top panel). (b) Thermal melt of Cy1 (blue) and Cy1 D391N (pink) monitored by circular dichroism at 222 nm. (c) Dynamics within Cy1 (blue) and D391N (pink) as determined by Hartman-Hahn R_ex_. (d) Population of minor conformers in Cy1 and D391N. A dynamic Cy1 engages with a loaded partner with an allosteric response (left cartoon) whereas a rigidified Cy1 is severely impeded (right cartoon).

## CONCLUSION

Overall, our studies establish global structural dynamics as universal sensors of molecular events and highlight challenges in interpreting outcomes of mutagenesis in dynamic proteins. Thus, structural fluctuations within a protein enable sensing of post-translational modifications within partners and can promote interactions together with remodelling of distant sites. Here, a point-site mutation cannot be interpreted solely through local effects, e.g. by considering the position of the mutation in a binding site (SI Discussion and SI Fig. 10). Instead, the mutation throws sand into intricate, concerted dynamics such that it affects binding events on remote sites, thus confounding interpretation through local criteria. As a corollary, our results indicate that intra-domain dynamics in Cy or C domains should be preserved in engineered NRPSs as they enable to remodel the inter-domain NRPS dynamics by promoting engagement with loaded partners and preventing unproductive interactions with holo domains (SI Discussion). The conventional wisdom is that structure is function, and our results show that function includes structural fluctuations. As structures become easier to determine, it is time to establish methods and models to routinely integrate dynamics to molecular mechanisms.

## METHODS

### Chemicals and reagents

Routine chemicals used for protein expression and purifications were purchased from Sigma, Fisher Scientific, VWR, Research Products International (RPI), or MP Biomedicals. NMR isotopes were purchased from Sigma (glucose and D_2_O for 4 samples used for NOESY experiments) or Cambridge Isotope Laboratories (all other samples). Salicylate, ATP, and DTT were purchased from Sigma, GoldBio, and VWR, respectively. Unless otherwise noted, *E. coli* strains for expression were purchased from Novagen. *E. coli* ΔEntD cell lines used in the expression of T1 were courtesy of Drs. Chalut and Guilhot (CNRS, Toulouse, France). Cloning and mutagenesis enzymes and primers were purchased from New England BioLabs (NEB) and Integrated DNA Technologies (IDT), respectively. The adenylation domain YbtE and thioesterase SrfAD were prepared as described in,^28^ with vectors provided by the C. T. Walsh lab (previously at Harvard Medical School). The precursors to prepare stereospecific ^1^H-^13^C-Me samples were a gift from Dr. Haribabu Arthanari (Dana-Farber Cancer Institute).

### Protein expression, purification, and *in situ* reactions

Holo-T1^33^ and Cy1^34^ were produced as previously described with variations in labelling schemes detailed in the SI Methods. The D391N variant was obtained through mutagenesis using PCR guided site-directed mutagenesis with primers 5’-gtctggataaatcatctggcgttcgagcatcacggcgaggtc-3’ (forward primer) and 5’-cagatgatttatccagacctgcggcgtttgcgagatgccccattc-3’ (reverse primer). Expression and purification protocols followed those used for wild-type Cy1 (SI Methods). Incomplete factorial design^35^ revealed conditions to optimize Cy1 stability. Following purification, Cy1 is stored in 20 mM sodium phosphate, 100 mM NaCl, 1 mM ethylenediamine tetraacetate (EDTA), 5 mM dithiothreitol (DTT), pH 8 at 4 °C. For deuterated samples, Cy1 was incubated at room temperature for 4 to 5 days in 50 mM Tris, 10 mM NaCl, 1 mM EDTA, 10 mM DTT, pH 8.5 to accelerate back-exchange of solvent-protected amides. The buffer for all NMR experiments, except for *in situ* experiments (*vide infra*), was 20 mM sodium phosphate, 10 mM NaCl, 1 mM EDTA, 5 mM DTT, 0.05 % w/v sodium azide, pH 7.0 at 25°C. D_2_O was added at 10 % v/v and sodium trimethylsilylpropanesulfonate (DSS) was added to reach defined concentrations in the range 0.2-0.5 mM (DSS protons). Cy1 is sensitive to oxidation and all buffers were freshly prepared with fresh DTT and degassed. NMR samples were degassed in Shigemi tubes, bubbled with argon, and sealed with glass plunger and parafilm.

Complexes of Cy1 or D391N with holo-T1 or loaded-T1 were prepared by co-concentration of individual stocks (prepared in identical buffers) to reach the desired concentrations. Here, 1D-NMR isotope-edited spectra were recorded for isolated proteins and for proteins in complexes to calculate the final experimental concentrations (details in SI Methods) by scaling concentrations initially determined using U.V absorption, with absorbance coefficients (ε) of 20,970 and 88,265 M^-1^ cm^-1^ for T1 and Cy1, respectively.

Holo-T1 was loaded with salicylate with a protocol modified from that of Goodrich et al.^33^ with the buffer 50 mM ACES, 10 mM NaCl, 2 mM MgCl_2_, 1 mM TCEP, pH 7 at room temperature. All NMR samples used in the *in situ* assay contained 10% D_2_O and DSS (0.2 mM DSS protons) for referencing. Loaded-T1 was presented four times to wild-type Cy1 and once to D391N. In each case, the following controls were performed. 2D-HN-TROSY^36^ were recorded for holo-T1, Cy1, and for holo-T1 with Cy1. 2D-IDIS-TROSYs^30^ were recorded for the latter, as well as after addition of salicylate to 1 mM and ATP to 2 mM. Loading of holo-T1 with salicylate were initiated by addition of the adenylation domain YbtE and monitored *in situ* through the emergence of signals characteristic of loaded-T1^28^ in HN-TROSY experiments (Figure 3, SI Figures 5 and 6). As discussed in the SI Discussion, Cy1 apparently accelerates hydrolysis of the thioester bond tethering salicylate to T1, suggesting that the reported hydrolase activity of C-domains^37^ extends to cyclization domains. We did not focus on this aspect other than varying the concentrations of YbtE, T1, and Cy1 such that we could detect Cy1 signals with sufficient signal-to-noise whilst controlling the amount of loaded T1. Depletion of ATP was monitored by 1D NMR and the integrity of loaded T1 was monitored by 2D HN-TROSYs. ATP needed to be replenished around 3 days after T1 was loaded to a stable amount. A first sample was prepared for quantifying the population of the minor conformer of Cy1 using holo-^1^H-^15^N-^12^C T1 and ^2^H-^15^N-^13^C Cy1 (Fig. 3 and SI Figs. 5, 6, and 9). Here, a 3D-HNCO was recorded for free Cy1 as well as before loading holo-T1 to verify that no spectroscopic changes occurred when it is presented to Cy1 (SI Fig. 6). T1 was then loaded to 58 % *in situ* and a second HNCO was immediately recorded. A second sample was prepared to help assign signals of the minor conformer of Cy1 using HNCO and HNCA recorded with ^2^H-^15^N-^12^C T1 loaded to 60% and ^1^H-Me-ILV-^2^H-^15^N-^13^C Cy1. To correlate the population of loaded T1 with that of the minor conformer of Cy1 (Fig. 3e), a series of 190 time-shared HN-TROSY/HC-HSQCs (each 7 minutes long) were collected at 5 nM YbtE, 50 nM YbtE, and 200 nM YbtE. Spectra were then summed in groups of 15, starting from the first spectrum recorded at 5 nM, with the next group starting from the third spectrum, and the last group containing the last 15 spectra recorded at 200 nM YbtE. Here, T1 was loaded to 80% in the final spectra recorded at 50 nM YbtE, with no additional loading observed at 200 nM YbtE.

A 3D HNCO was recorded when D391N was presented with loaded-T1 using an *in situ* protocol, with controls first performed, and holo-T1 loaded upon addition of YbtE to 100 nM. D391N and T1 concentrations were similar to that used for WT HNCO and HNCA (SI Methods). This time, T1 was loaded to 77%. We did not reduce the concentration of YbtE to reach the same conversion rate as for wild-type as the larger population of loaded-T1 strengthened our observation that D391N does not respond to loaded-T1 as efficiently as the wild-type Cy1 does. It is tempting to extrapolate that the mutation impacts the hydrolase activity of Cy1, but we had not conducted controls with free holo-T1 during that set of experiments, and future studies should be performed to validate this observation (SI Discussion).

To demonstrate that the minor conformer of Cy1 (WT) does not result from the build-up of AMP or Sal-AMP during the *in situ* reaction, we added the thioesterase SrfAD to 500 nM and verified that T1 was restored to its holo form. A 3D-HNCO was then recorded to confirm that the signals of the minor conformer disappeared (SI Fig. 6), thus confirming that they reflected specifically a response of Cy1 to salicylate attached to T1 through a thioester bond.

### NMR data acquisition and processing

All experiments were conducted at 25 °C on a 600 MHz AVANCEIII Bruker spectrometer (Hopkins School of Medicine) equipped with a QCI cryoprobe, an 800 MHz Varian Unity+ spectrometer equipped with a chiliprobe (Hopkins Arts and Science), and a 950 MHz AVANCEIII Bruker spectrometer (University of Maryland School of Medicine) equipped with a TCI cryoprobe. *In situ* experiments and Hahn-Echo experiments were recorded at 600 MHz, experiments to assign resonances were recorded at 600 and 800 MHz, distance constraints were measured at 600, 800, and 950 MHz, and relaxation dispersion profiles were recorded at 600 MHz and 950 MHz (*vide infra*). A list of experiments featuring acquisition parameters, samples, and fields is provided in the SI Methods. All experiments used for Cy1 used TROSY^36^ to minimize losses due to relaxation. All spectra were processed with NMRPipe^38^ and analysed with CARA^39^ (assignments) or Sparky^40^ (chemical shift perturbations).

### Assignment of backbone resonances

Details of acquisition parameters, experiments, and samples are provided in the SI Methods. Notable strategies to overcome spectral crowding included using the TROSY-hNCAnH^41^ and TROSY-hNcaNH^42^ experiments as well as calculating 4D covariance correlation maps with non-uniformly sampled HNCO, HN(CA)CO, HN(CO)CA, HNCA, HN(COCA)CB, and HN(CA)CB. In the end, 93% of Cy1 backbone resonances were assigned. The mutant D391N was assigned through transposition of assignments from wild-type Cy1 using HNCA and HNCO. 82% of the mutant backbone resonances were assigned. Assignments of T1 were reported before.^33^

### Assignment of side-chain resonances

The majority of Cy1 methyl resonances were assigned using covariance maps calculated through HMCM(CG)CB, HMCM(CGCB)CA, HNCA, and HN(CA)CB spectra^34^ collected for a ^1^H-Me-ILV-^2^H-^15^N-^13^C sample. HCCONH^43^ experiments were recorded on the same sample together with an HCCH-TOCSY. 98 % of Cy1 methyls (ILV) were assigned. 42% of aromatic (tryptophan, phenylalanine, and tyrosine) sidechain proton signals were assigned following initial structure calculations, using NOESY spectra and H^α^ assignments from MQ-HACACO^44^ as check points.

### NMR constraints

Torsion angle constraints were obtained with the program TALOS+.^45^ The majority of the distance constraints were determined with a time-shared 3D HN-TROSY/HC-HSQC-NOESY^46^ with a mixing time (τ_m_) of 40 ms collected at 950 MHz on a ^1^H^13^C-δ1I-δ2L-γ2V-^2^H-^15^N sample.^47^ A second spectrum was collected for long-range constraints (τ_m_ 150 ms). Assignments were facilitated through a non-uniformly sampled 4D HN-TROSY/HC-HSQC-NOESY-HN-TROSY/HC-HSQC experiment^48,49^ (τ_m_ 200 ms), recorded at 800 MHz on a ^1^H^13^C-δI-LV-^2^H-^15^N sample that also contained protonated tyrosine and phenylalanine.^50^ A 3D NOESY-HN-TROSY/HC-HSQC was also recorded to provide constraints with aromatic moieties. Other NOESYs were collected on Cy1 samples during method developments, and distance constraints were retrieved as described in the next section and the SI Methods.

^1^H-^15^N residual dipolar couplings were obtained by recording TROSY and ^15^N decoupled 3D HNCOs in an interleaved manner in both isotropic (NMR buffer) and aligned media. Alignment was achieved using 12 mg/mL of Pf1 phage (deuterium quadrupolar splitting of 11.4 Hz). Residues displaying relaxation dispersion or with ^15^N order parameters smaller than 0.7 (obtained from TALOS+^45^) were excluded from the restraint list.

### Structure calculation

Details in the SI Methods. Structures were calculated using CYANA 3.98^51,52^ and refined using CNS.^53^ 810 dihedral angle restraints were prepared using a consensus between TALOS+^54^ and CSI 3.0.^55^ 293 hydrogen bond constraints were introduced based on secondary structure prediction through TALOS+ and NOE patterns. A structural model calculated with RASREC CS-Rosetta ^56^ (SI Methods) was used to provide hydrogen bond and dihedral angle constraints for unassigned residues or residues whose signals were too weak to provide constraints; 7 residues in α4 and 5 in β11. First, a structural ensemble (root-mean-square deviation (RMSD) of 1.96 Å over residues in secondary structured elements) was calculated using 1187 accurate distances obtained from time-shared NOESY spectra collected on a sample with stereospecific protonation of Leu and Val methyls at δ2 and γ2, respectively, and with a mixing time of 40 ms, i.e. with minimal spin diffusion. A representative conformer was then identified from the resulting structural ensemble. To include long-range constraints, an additional 1,002 distance restraints were selected from a set of NOESY spectra at longer mixing times (excluding those at 40 ms, see SI Methods) and binned into groups with upper limits at 3, 5, 7, and 9 Å such that they were in agreement with the initial representative model. 176 residual dipolar couplings were then incorporated, excluding those of residues showing dispersion or possessing low order parameters (estimated through TALOS+). The deposited bundle features no violations larger than 0.5 Å and exhibits an RMSD of 1.2 Å for residues in secondary structured elements (SI Table S1).

### Experiments to determine protein dynamics

Relaxation dispersion experiments at 600 and 950 MHz were recorded with TROSY and relaxation compensation of constant-time Carr-Purcell-Meiboom-Gill pulse sequences,^57,58^ using a 600 μM sample of Cy1 with ^1^H^13^C-δI-δ2Lγ2V-^2^H-^15^N labelling. The complete set of relaxation dispersions were recorded in an interleaved manner such that all CPMG frequencies were sampled before indirect nitrogen dimensions were encoded. The CPMG frequencies were acquired in random order. Intensities were estimated through line-shape fitting with the nLinLS module of NMRPipe.^38^ The relaxation dispersion profiles were fitted using ChemEx,^59^ which integrates the Bloch-McConnell equation over the NMR pulse sequence. The fit was run in three consecutive steps. First, a subset of isolated signals showing dispersions and good signal-to-noise were selected and fit together to obtain an estimate of the population and exchange rate for a two-site model. Second, all profiles were fit with exchange rates and populations fixed to the values obtained in the first step. Third, residues for which the change in chemical shift was smaller than the estimated uncertainty were kept fixed at a value of 0.0 during a final fit. Of 298 residues integrated, 178 could be fit. The profiles and their fits were subject to further statistical analysis following data fitting. Dispersion was considered significant only if the following two criteria were fulfilled. 1) Their data could not be fitted by a single average value,^60,61^ as measured by the following χ^2^:

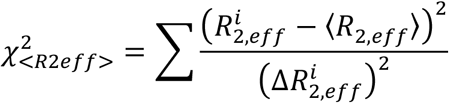

where 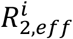 denotes the effective transverse relaxation rate for a given CPMG frequency n_i_,^57^ <> denotes the mean value over all frequencies, and 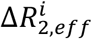 denotes the uncertainty as described in ^60^. 2) The magnitude of the fitted dispersion had to exceed the average residual of the fit. 52 residues were used for quantitative comparisons of chemical shifts based on low χ^2^ at both 600 MHz and 950 MHz, another 74 residues showed unambiguous dispersion, and 5 residues clearly demonstrated departure from the global fit.

Structural fluctuations over a wider μs-10’s of ms timescale were detected through the Hahn-Echo method with a 600 μM sample of Cy1 WT (CDN) and a 122 μM sample of Cy1 D391N (CDN) using the analysis described in ^32^ and a pulse sequence modified to account for ^13^C labelling. See SI Methods for more details.

### Calculation of Chemical Shift Perturbations (CSPs, Δδ)

For all CSPs, samples were prepared in an identical buffer, spectra were referenced through DSS,^62^ and temperatures were calibrated. CSPs were calculated through the relation:^63^

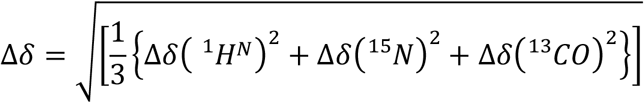

where δ is the chemical shift difference in ppm for a given nucleus between the two sets of signals that are compared (e.g. Cy1 WT and Cy1 D391N or Cy1 major conformer and Cy1 minor conformer). For figures where the CSPs are mapped onto the medoid structure of Cy1, residues are colored according to their CSP value with a gradient ranging from grey to red, where red indicates a CSP that is at or above two standard-deviations from the median value of CSPs. See SI Methods for more details.

### Cy1 Tunnel Calculations

All conformers from the Cy1 NMR ensemble were analysed using the CAVER 3.0.3 Pymol plugin.^64^ For each conformer, tunnels were generated by optimizing parameters such that a tunnel connecting the putative donor and acceptor thiolation domain entrances in the structure were found (details in SI Methods). To identify closed tunnels, where donor or acceptor entrances were inaccessible, the probe radius used in the calculation was decreased and the obstructed tunnels were identified from a set of unrealistic tunnels by comparison with open tunnels in other conformations.

### Cy1 WT and D391N Thermal Stability

Unlabelled wild-type Cy1 (WT) and D391N samples were spectroscopically characterized using circular dichroism (CD). Cy1 WT and Cy1 D391N at concentrations of 0.035 and 0.028 mg/mL, respectively, were thermally unfolded in NMR buffer (20 mM sodium phosphate, 10 mM NaCl, 1 mM EDTA, pH 7) with 5 µM TCEP and 0.0025% w/v NaN_3_. For CD measurements, a wavelength of 222 nm was monitored over the course of thermal unfolding. Both Cy1 WT and D391N exhibited an irreversible, two-state unfolding transition and were fit to a two-state model for determination of melting temperatures (see SI Methods).

### Statistics and Error Analysis

See also details of experimental procedures. The experimental error bars in the relaxation dispersion profiles (Fig. 3) are calculated as described in ^60^ following integration through nLinLS with NMRPipe.^65^ The error on reported chemical shifts (SI Fig. 4, 6, and 7) were estimated using the covariance matrix approach.^66^ When determining the percentage of loaded-T1 and minor peaks of Cy1 as T1 was loaded *in situ* (Fig. 3e), populations were calculated as:

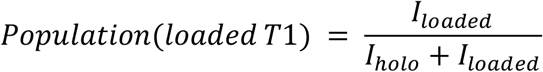

and

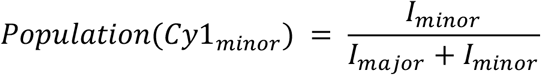

The values reported are means over two signals (T1) and six signals (Cy1) detected in time-shared experiments and the error bars were calculated via error propagation where the noise of the associated spectrum is the uncertainty of measurement of each signal intensity. In other experiments, loading of T1 was quantified through five signals resolved in 2D IDIS-TROSY^30^ data. Means were calculated and the error bars in SI Figs. 5 and 9 represent standard deviations to the mean. The populations of Cy1 minor residues are reported as a scatter plot in Fig. 4d, with a box-and-whisker in which the central line denotes the median and the edges the 25^th^ and 75^th^ percentiles. The limits of the whiskers identify populations with more than 1.5 times the interquartile range. Only 5 signals were detected for the minor conformer of D391N, and they could not be assigned. For all other residues, the noise amplitude defines a threshold for detection that was used to calculate the population, which lead to a 10-fold reduction in population when compared to wild-type Cy1. For Cy1 thermal melt analyses (Fig. 4b and SI Fig. 8), the error on the signals was computed using the standard deviation of the signal averaged over one second on the CD instrument. Error bars in the determination of Hartmann-Hahn R_ex_ profiles for Cy1 WT and D391N (Fig. 4c) were determined using Monte Carlo error estimates over 300 iterations, with the noise used as the variance for the distribution.

## Data Availability

The data that support the findings of this study are available from the corresponding author upon reasonable request. NMR chemical shifts have been deposited in the Biological Magnetic Resonance Data Bank under the following accession codes: 30943 for wild-type Cy1 and 51105 for mutant D391N. The structure of Cy1 has been deposited to the Protein Data Bank (PDB) under accession code 7RY6.

## Acknowledgements

We thank Drs A. Majumdar (Johns Hopkins Biomolecular NMR Center), Katie Tripp (Hopkins Center for Molecular Biophysics), David Weber and Kristen Varney (University of Maryland School of Medicine), and Bob Cole (Hopkins Mass Spectrometry and Proteomics Facility) for their assistance. This work was supported by NIH grant GM104257 to D.P.F.; N.S. acknowledges funding from NIAID grant 5R01Al143997 and NIGMS grant 5R35GM125034; D.P.D acknowledges funding from NIH grant 1R15GM123425-01.

## Author Contributions

S.H.M., A.K.K., K.A.M. and D.P.F. designed research and wrote the manuscript. S.H.M. assigned the resonances of Cy1 and its distance constraints, with help from D.D.’s EpoB crystal structure model, determined the initial Cy1 NMR structure model, performed *in situ* reactions with A.K.K. and K.A.M., and collected data for Hahn-Echo and relaxation dispersion for Cy1. In addition, A.K. determined the structure of Cy1 and analysed RDCs. K.A.M. collected (assisted by S.H.M. and A.K.K.) and analysed all data for D391N, collected and analysed CD data for Cy1, performed Cy1 tunnel calculations, and determined minor Cy1 populations. S.N. and N.S. calculated a RASREC model of Cy1’s C-terminal domain, and G.B. analysed the relaxation dispersion data.

## Supporting Information

is available separately.

