## Supporting Information for "Global Dynamics as Communication Sensors in Peptide Synthetase Cyclization Domains"

**THIS WORK HAS NOT BEEN PEER-REVIEWED.**

† Present address: Reference Standard Laboratory, United States Pharmacopeial Convention, 12601 Twinbrook Pkwy, Rockville, MD, 20852, USA

### SI Table:

**Table S1. NMR and structure refinement statistics for Cy1[a]**

| | |
|---|---|
| **Completeness of resonance assignments[b]** | |
| **Backbone amide resonances** | **93%** |
| **Methyl resonances** | **98%** |
| **Conformationally-restricting restraints[c]** | |
| Total NOEs | 2189 |
| Short ($\|i - j\| \leq 1$) | 684 |
| Medium range ($1 < \|i - j\| < 5$) | 690 |
| Long range ($\|i - j\| \geq 5$) | 815 |
| Hydrogen bond restraints | 293 |
| Total Dihedral angles | 810 |
| $\phi$ | 395 |
| $\psi$ | 415 |
| $^1D_{NH}$ RDC restraints in Pf1 phage alignment medium | 176 |
| Total restraints | 3468 |
| Restraints per residue | 7.9 |
| **Structure Statistics** | |
| **Residual restraint violations[d,e]** | |
| RMS distance violation / restraint (Å) | 0.02 |
| Maximum distance violation (Å) | 0.44 |
| RMS dihedral violation / restraint (°) | 1.28 |
| Maximum dihedral angle violation (°) | 16.00 |
| **RMSD from average coordinates (Å)** | |
| All Backbone atoms (selected/ordered/all)[f,g] | 1.2/1.5/2.4 |

| | |
|---|---|
| All Heavy atoms (selected/ordered/all)[f,g] | 1.8/2.1/2.9 |
| **Deviation from idealized geometry** | |
| RMSD Bond lengths (Å) | 0.02 |
| RMSD Bond angles (°) | 1.2 |
| **Molprobity Ramachandran plot** | |
| Most favored regions (%) (selected/all)[f] | 99.0/94.4 |
| Allowed regions (%) (selected/al)[f] | 0.9/4.7 |
| Disallowed regions (%) (selected/all)[f] | 0.1/0.8 |
| MolProbity[1] clashscore (mean/Z) | 7.74/0.20 |

[a] Structural statistics were computed for the ensemble of 20 deposited structures of Cy1 using PSVS 1.5.[2]

[b] Backbone amides of N-terminus and C-terminal hexahistidine tag are excluded. Observable methyl resonances consistent with selective $CH_3$ labeling in otherwise deuterated background are included: Ile (δ1), Leu (δ1,2), Val (γ1,2).

[c] There are 439 residues with conformationally restricting constraints.

[d] Analyzed for all residues 1-453.

[e] Average distance constraints were calculated using the sum of $r^{-6}$.

[f] Selected residues include all those in regular secondary structured regions viz., α-helices and β-strands: 22-29,42-48,53-65,72-75,79-82,91-95,101-117,130-134,140-146,153-168,181-209,230-237,239-252,256-270,277-284,301-308,315-329,338-347,356-360,390-397,400-408,414-433

[g] Ordered residues include those whose sum of phi and psi order parameters > 1.8: 10-37,41-47,52-86,90-124,126-147,149-151,153-169,171-297,301-354,356-365,368-380,388-405,407-443

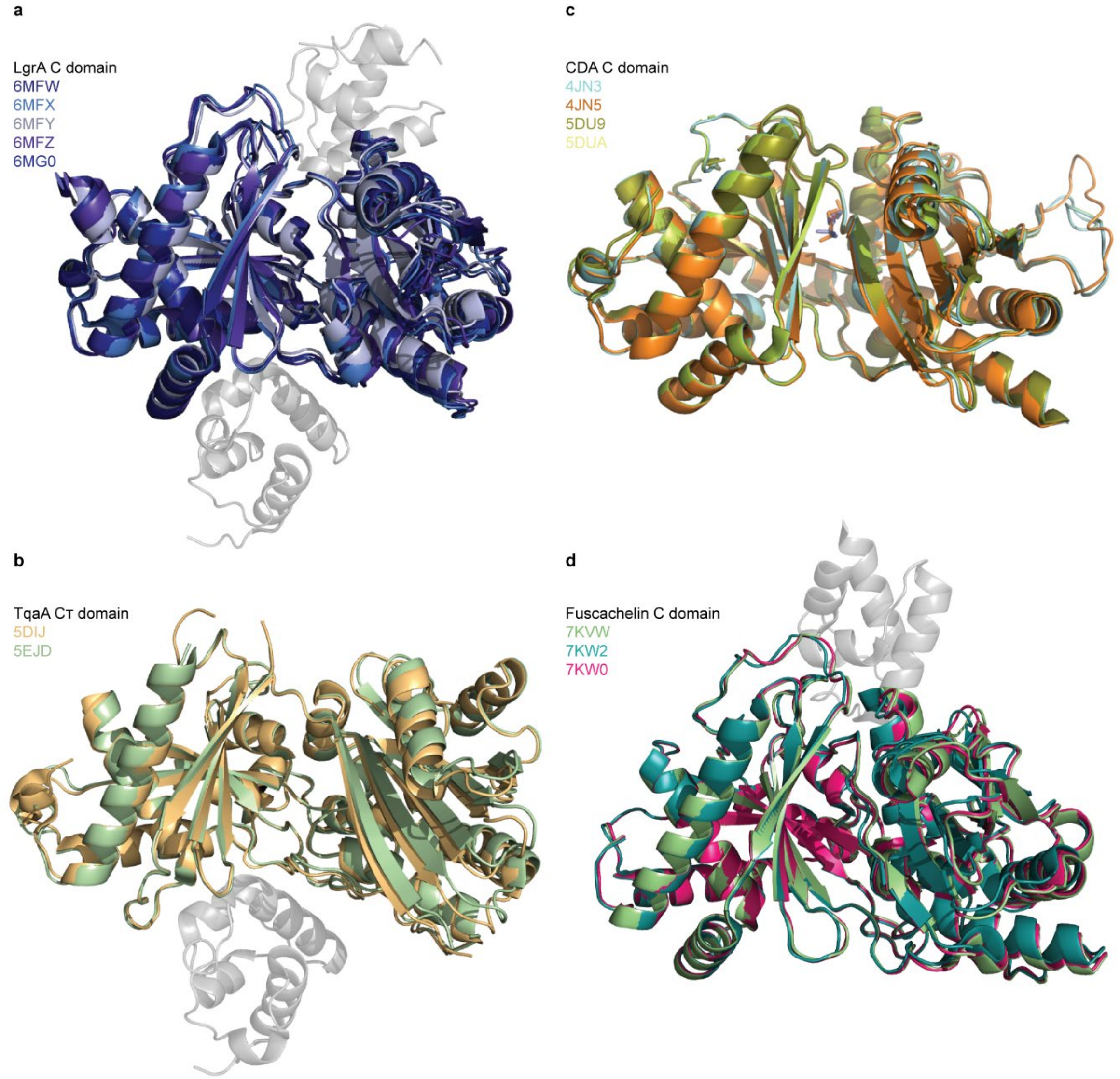

**SI Figure 1. Comparison of C-domain crystal structures in the presence and absence of thiolation domains and substrates.** **a**, LgrA C domain[3] with a docked donor thiolation domain (PDB: 6MFW, 6MFX, 6MFY), docked donor and acceptor thiolation domains (6MFZ), and in the absence of thiolation domains (6MG0). Donor (bottom) and acceptor domains (top) from 6MFZ are shown in grey. **b**, TqaA C-terminal domain[4] with (5EJD) and without (5DIJ) a docked donor T domain (in grey). **c**, CDA-C1 domain without (4JN3, 4JN5)[5] and with substrate analogues (PDB: 5DU9, 5DUA)[6]. **d**, Fuscachelin C domain[7] with a holo thiolation domain at the acceptor site with the phosphopantetheine arm outside the C-domain tunnel (PDB: 7KVW), within the C-domain tunnel (PDB: 7KW2), and with a substrate-loaded thiolation domain harboring an arm and substrate within the tunnel (PDB: 7KW0). The donor thiolation domain from 7KVW is shown in grey. Alignments performed in PyMOL 2.0.7 using the *align* command on all $C^{\alpha}$ atoms in the C-terminal region (lobe on the left).

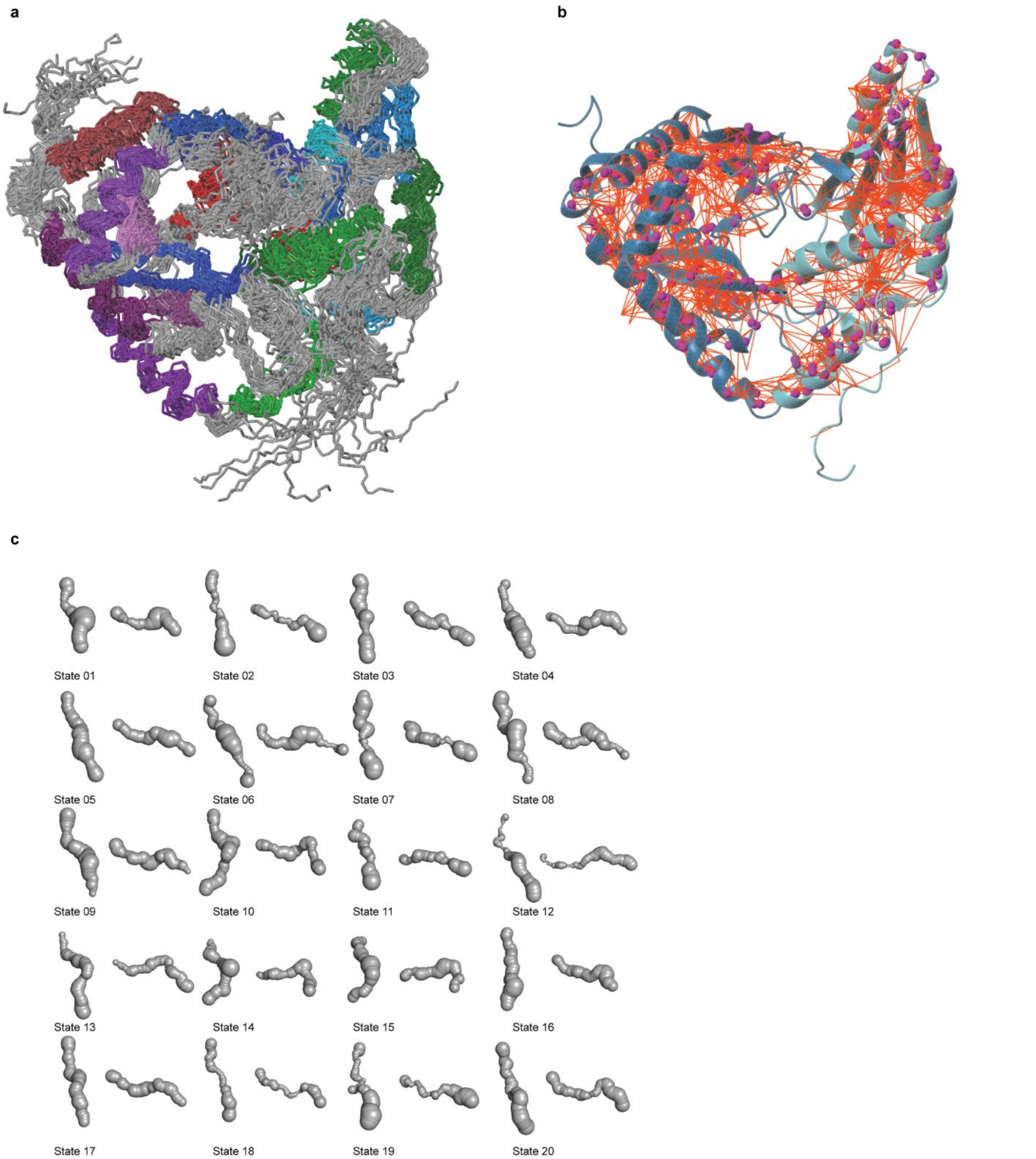

**SI Figure 2. Ybt Cy1 NMR ensemble. a**, Ybt Cy1 NMR ensemble aligned on major secondary structured elements (colored). **b**, Distance (orange lines) and residual dipolar couplings (magenta spheres) restraints used in the Cy1 structure calculation. **c**, Tunnels between donor and acceptor sites calculated using CAVER. Each tunnel is displayed with a top view (as in Figs. 2, 3, and 4) and a side view (rotation of 45 along X and 90 along Y). Structural heterogeneity within the tunnel and at the acceptor and donor sites modulate the tunnel shape.

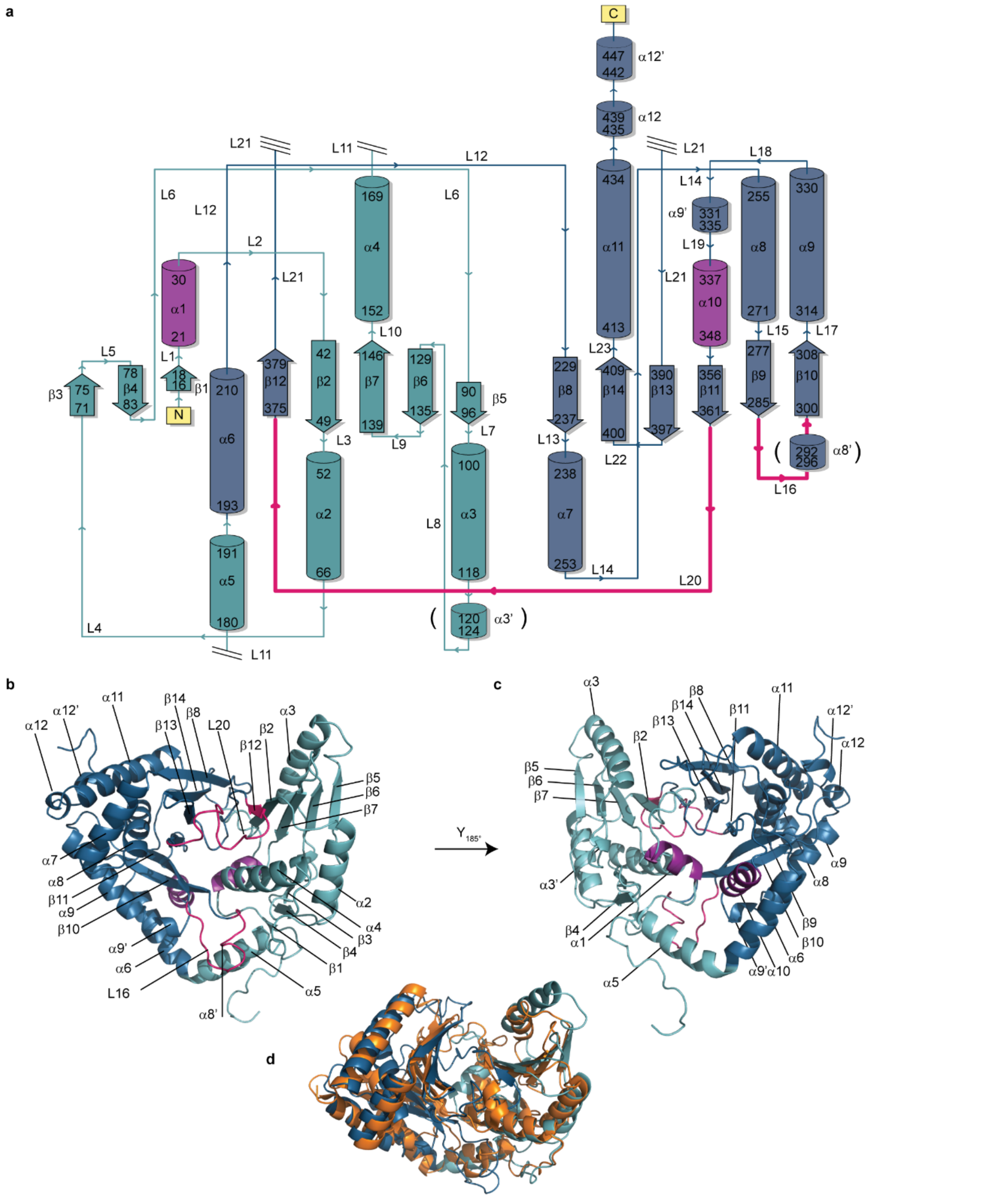

**SI Figure 03. Structural Features of Cy1.** **a**, Topology diagram (PDBsum). Primes denote structural elements present in Cy1 that are not always present in the C-domain family. Parentheses denote elements that are not always seen in the NMR ensemble. Hot pink: donor site. Purple: acceptor site **b** , **c**, Structural elements on the medoid conformer of the NMR ensemble. **d**, Alignment of Cy1 medoid structure with all members of the C-domain family ($C^{\alpha}$ carbons in secondary structures using the *super* command in PyMOL 2.0.7) reveals its conformation to resemble most that of LgrA[3] (6MFZ, orange).

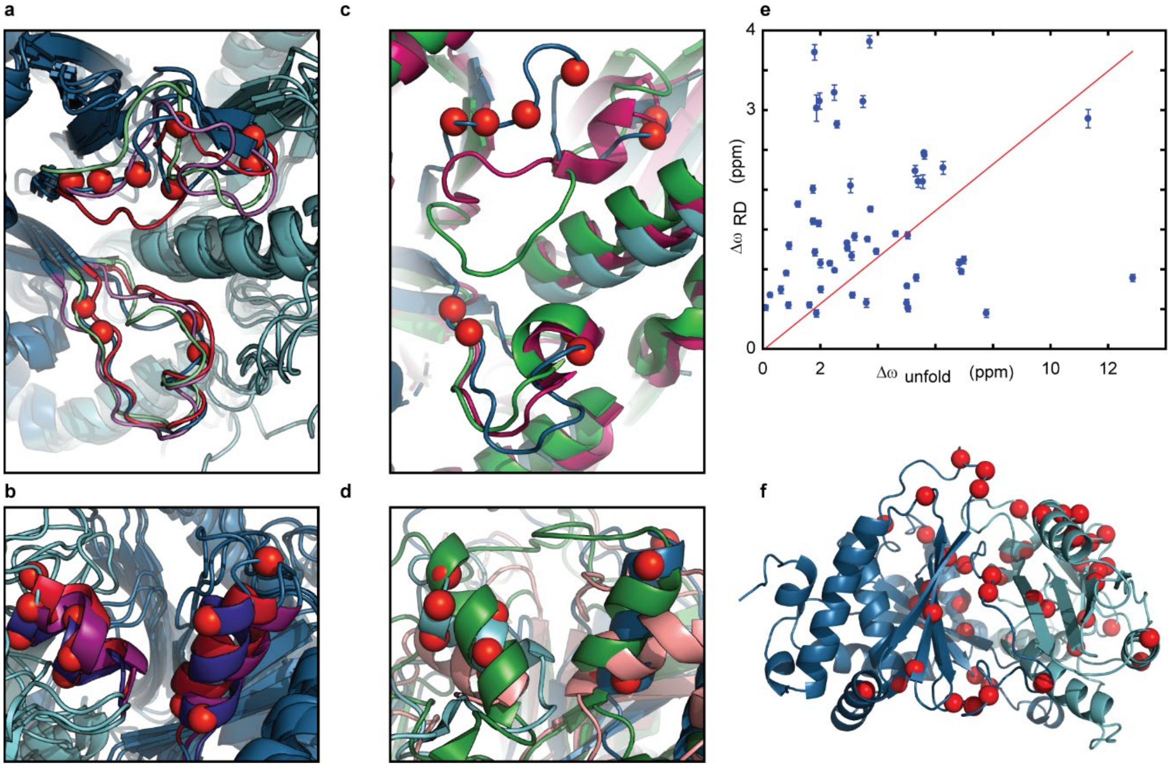

**SI Figure 4. Malleability of donor and acceptor thiolation domain binding sites in the C-domain family.** **a**, Donor thiolation domain site in Ybt Cy1 NMR conformers wherein L20 and L16 are colored differently for three conformers displaying open (green), closed (red), and half-open (blue) conformations, as identified by tunnels. **b**, Structural heterogeneity in the acceptor site of Cy1 NMR conformers wherein $\alpha$1 and $\alpha$10 have been colored according to conformers. **c**, Donor site in the BmdB Cy2 domain[8] (magenta), the EpoB Cy2 domain[9] (green), and our NMR medoid structure (blue). **d**, Acceptor site of the EpoB Cy domain (green), the GrsA E domain[10] (gold), and our NMR medoid structure (blue). In **a** – **d** and **f**, red spheres denote relaxation dispersion and hence fluctuating environments. The conformational changes captured by crystallography in **c** and **d** mirror the fluctuations determined by NMR shown in **a** and **b**. **e**, Absence of correlation between chemical shifts from relaxation dispersion and unfolding ($R^2 = -0.82$). **f**, Residues analyzed in **e** encompass the entire protein fold. Alignments were performed over $C^\alpha$ carbons in secondary structures in PyMOL 2.0.7 using either the *align* command (**a, b**) or the *super* command (**c, d**).

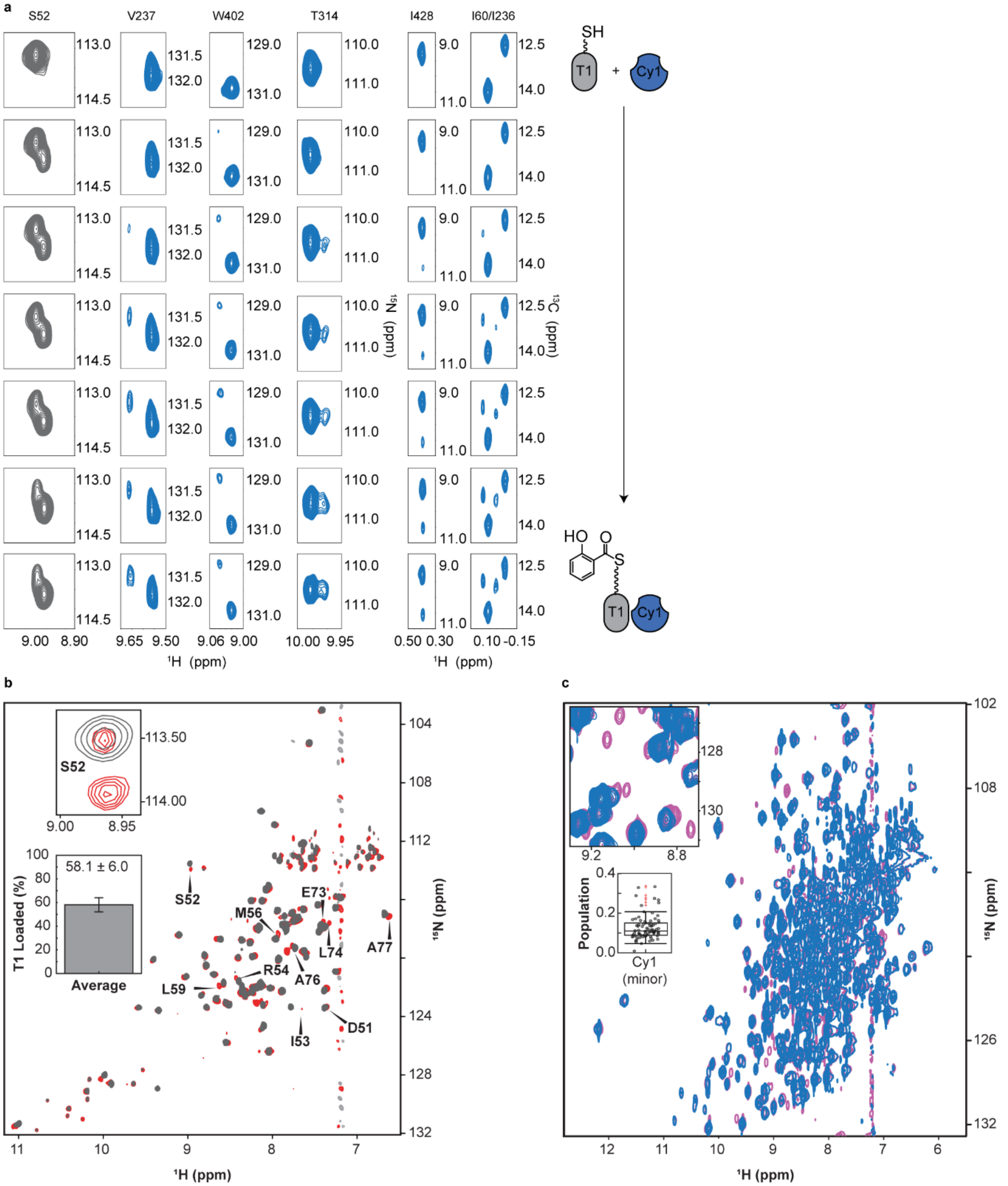

**SI Figure 5. *In situ* loading of T1 in the presence of Ybt Cy1. a**, Zooms of time-shared NMR spectra showing substrate loading of holo-T1 (grey, S52) with concomitant response of Cy1 (blue) over time. Cy1 response is shown through the amides of V237, W402, and T314 in addition to the methyl signals of I428 and I60/I236. **b**, When T1 is loaded to a steady state, the IDIS method enables to separate the spectra of $^{2}$H-$^{15}$N-T1 (**b**) from those of $^{2}$H-$^{15}$N-$^{13}$C Cy1 (**c**). In **b**, the IDIS-TROSY spectra of T1 show holo-T1 (grey) and salicylate-loaded T1 (red) when in presence of Cy1. The first inset shows signals of S52, the conserved site for phosphopantetheinylation. The bar plot in the second inset shows the average percentage of loaded-T1 observed in the IDIS-spectrum. The error bar indicates the standard deviation over five T1 residues used to estimate this average (see SI Methods). In **c**, the IDIS-TROSY spectrum of Cy1 shows Cy1 in complex with holo-T1 (blue) and with loaded-T1 (pink). The first inset shows an example region of the NMR spectrum where minor Cy1 peaks emerge in response to salicylate loaded T1. The second inset shows the population of these minor peaks as determined using a 3D HNCO NMR experiment (see SI Methods).

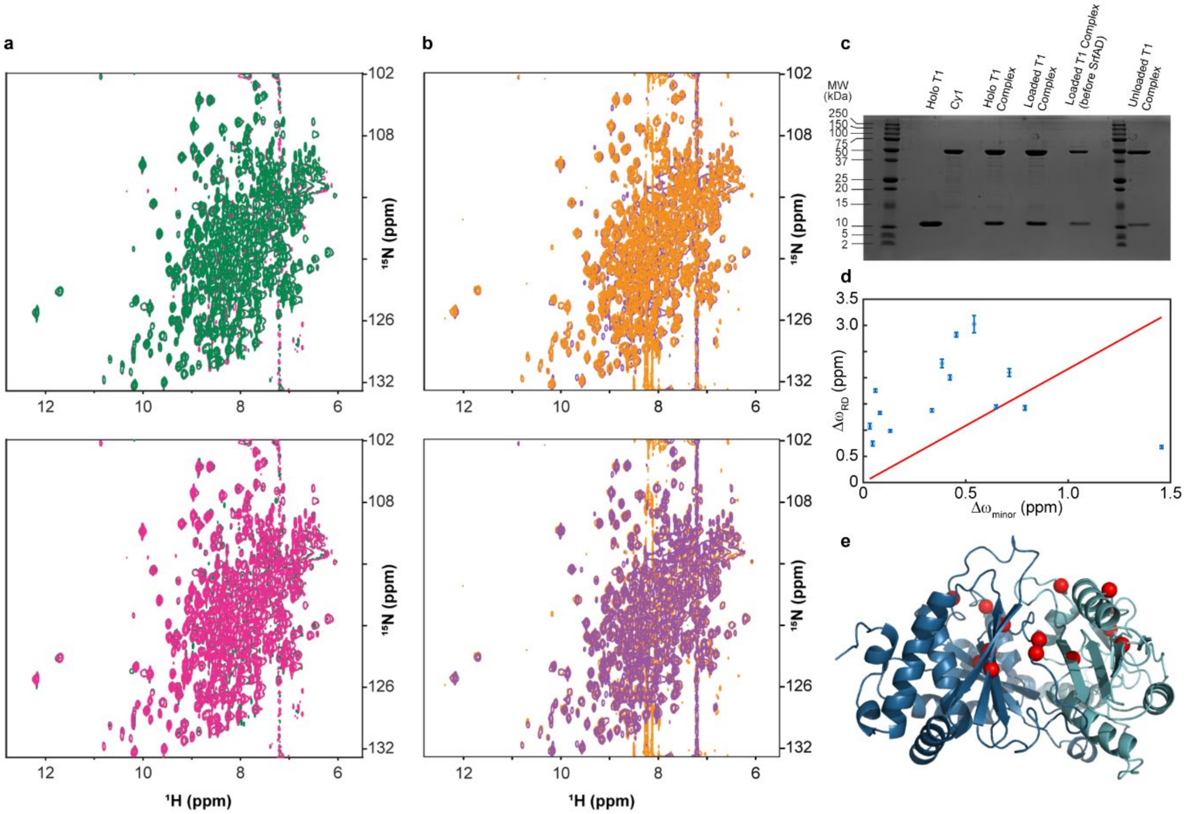

**SI Figure 6. Cy1 WT *in situ* controls. a**, Overlays of 2D HNCO projections of free Cy1 (green) and Cy1 in a complex with holo-T1 (pink). Top: free Cy1 in foreground. Bottom: free Cy1 in background. **b**, Overlay of 2D HNCO projections of Cy1 in complex with loaded-T1 (purple) and Cy1 in complex with unloaded-T1 following addition of the thioesterase, SrfAD (gold). **c**, SDS-PAGE gel of Cy1 *in situ* NMR samples before and during the *in situ* reaction and after addition of SrfAD. **d**, Comparison of changes in $^{15}N$ chemical shifts, Δω, obtained from relaxation dispersion (RD) with those obtained by comparing signals of Cy1 minor and major conformers upon addition of loaded-T1. The absence of correlation ($R^2$ = - 2.4) indicates that the allosteric response of Cy1 does not occur through simple conformational selection of a minor state in free Cy1. **e**, distribution of residues analyzed in **d**. In **d**, we excluded all residues with shifts that may be impacted through molecular contact. The residues were identified using the *within* command in Pymol with values of 2 Å, for the phosphopantetheine arm and T1, and 6 Å, for salicylate. The positions of the T-domain and phosphopantetheine arms were taken from the LgrA[3] system. These numbers refer to [11] and reflect the distances at which chemical shifts may change without conformational changes for Cy1 residues. The effect of salicylate was represented by a sphere of 6 Å at the center of salicylate as defined by 2N6Z[12]. Although only 14 residues could be analyzed in **d**, these residues nevertheless span the protein fold as shown in **e**.

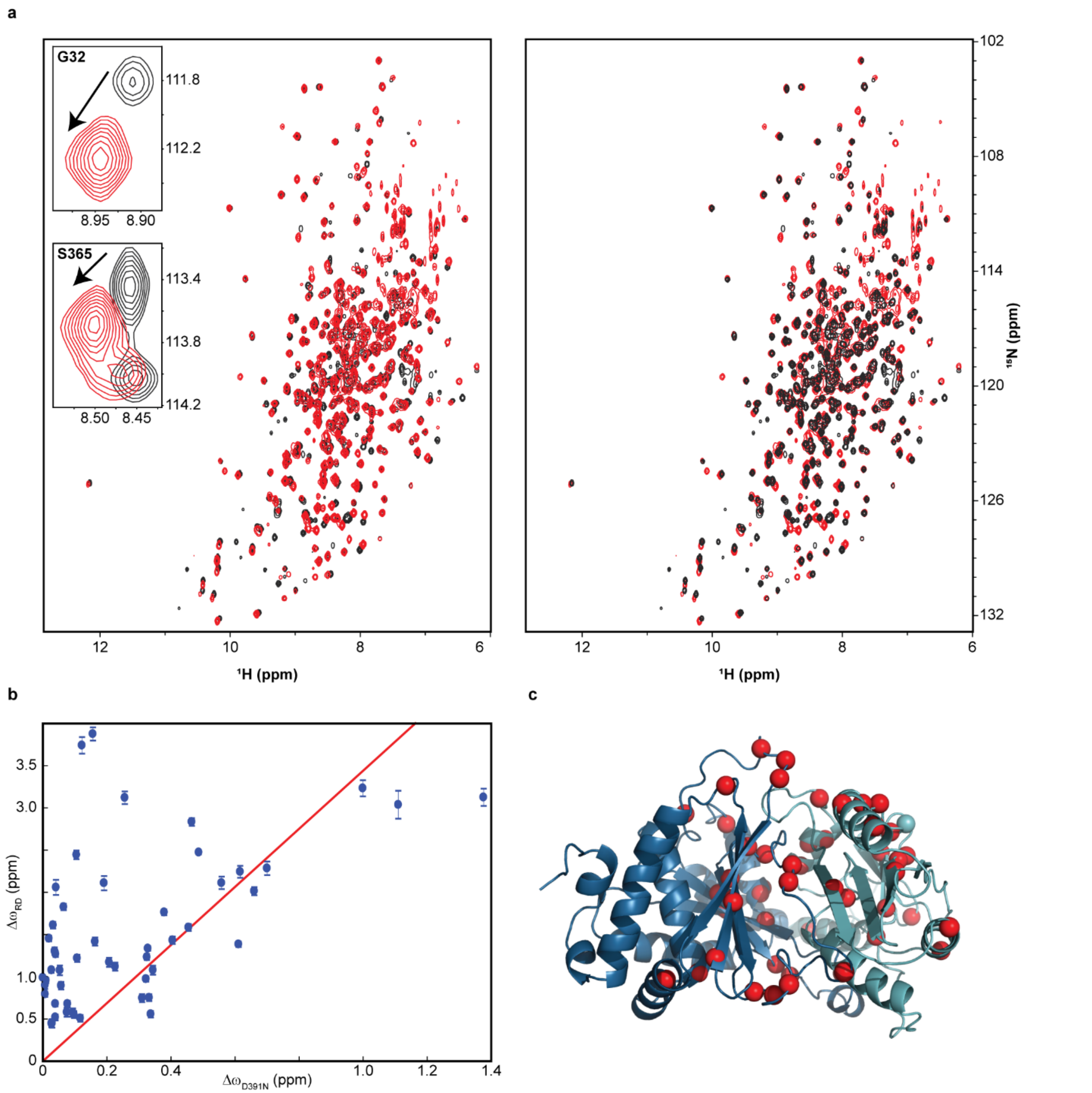

**SI Figure 7. Impact of the D391N mutation on the environment of Cy1 residues. a**, Overlay of 2D HN-TROSY of wild-type (black, background) and D391N Cy1 (red, foreground). Insets show chemical shift perturbations at the acceptor site (G32) and at the donor site (S365). On the right, the spectrum of wild-type Cy1 is in the foreground and that of D391N is in the background. The mutation impacts residues throughout the entire protein (Fig. 4). **b**, Comparison of changes in chemical shifts, $^{15}$N Δω, obtained from relaxation dispersion measured on wild-type Cy1 and obtained by comparing wild-type Cy1 with D391N. The absence of correlation ($R^2 = -0.52$) indicates that the mutation does not simply select for a conformer on the path between a major and minor conformation of Cy1. **c**, The residues analyzed in **b** (red spheres) encompass the fold of Cy1.

a

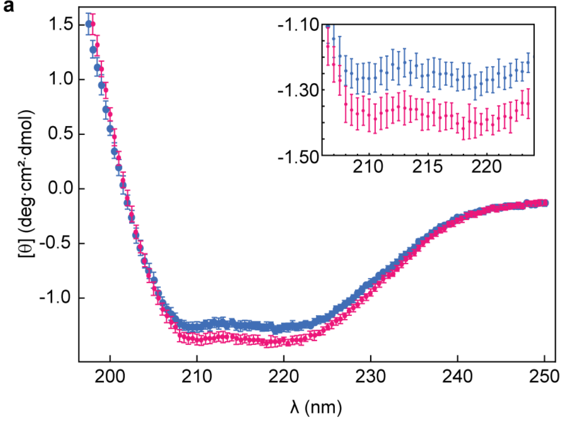

b

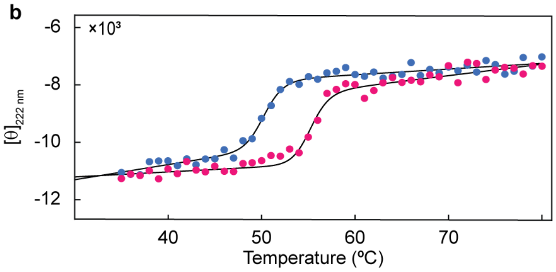

c

| | Cy1 WT | Cy1 D391N |
|---|---|---|
| $T_m^{app}$ (°C) | 50.32 ± 0.16 | 55.24 ± 0.20 |

d

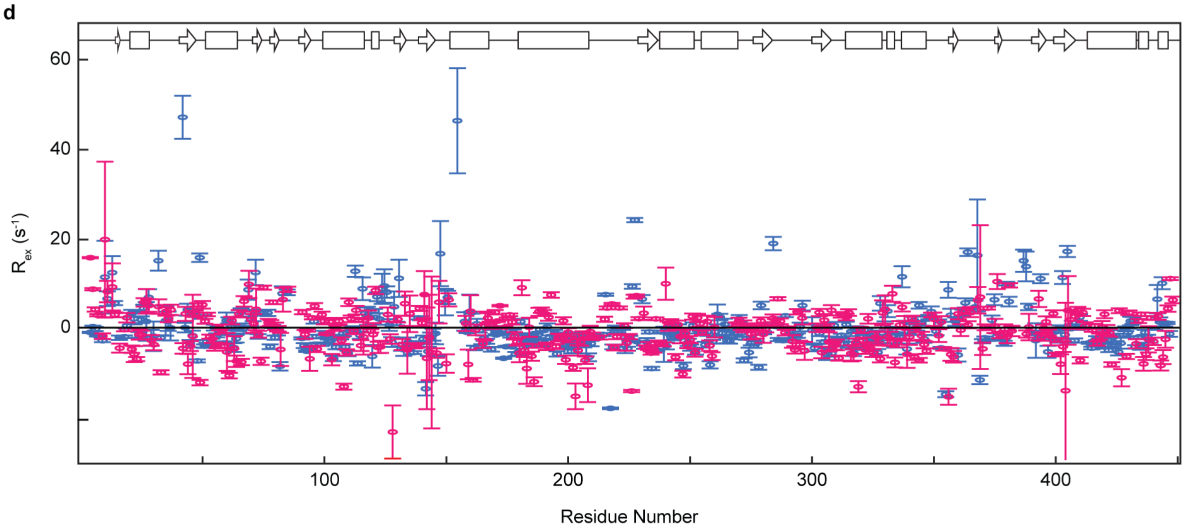

**SI Figure 8. Spectroscopic and dynamic analysis of Cy1 WT and D391N. a**, Circular dichroism (CD) spectra of Cy1 WT (blue) and D391N (pink). Inset shows spectra expanded over the range 205 to 224 nm. **b**, Thermal melt of Cy1 monitored by CD at 222 nm. Fitted melting curves are shown in black. **c**, Fitted Cy1 apparent melting temperatures ($T_m^{app}$) for CD data shown in **b** using the two-state unfolding equation described in SI Methods. **d**, Cy1 dynamics as determined by Hartmann-Hahn $R_{ex}$ profiles. This panel is the same as shown in Fig. 4c but enlarged to show detail. Cy1 secondary structure is shown at the top of the figure where beta sheets are denoted with arrows and helices with rectangles.

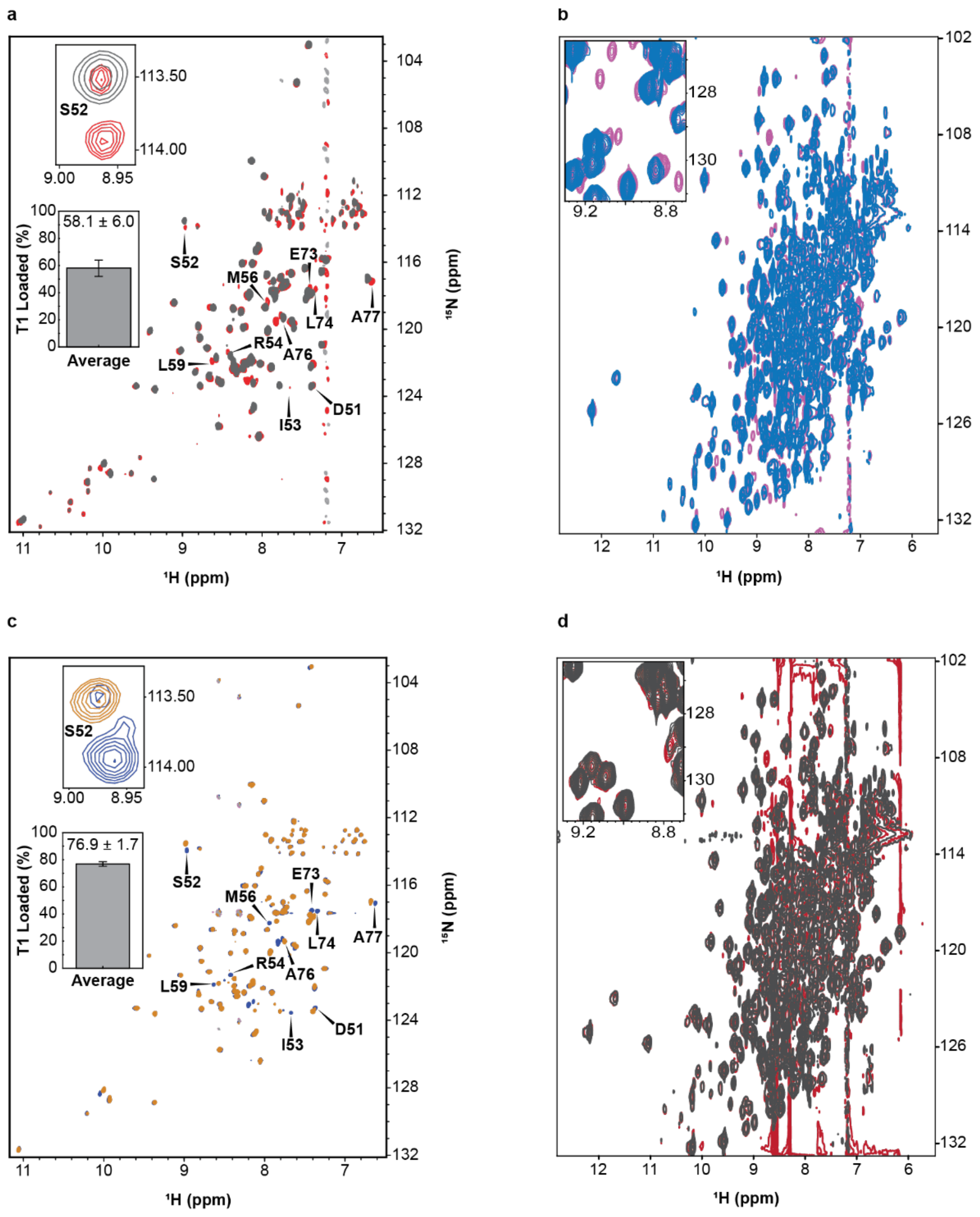

**SI Figure 9. *In situ* loading of T1 in the presence of wild-type Cy1 and D391N.** IDIS-TROSY spectra allow for simultaneous inspection of T1 and Cy1 as T1 is converted from holo to salicylate-loaded forms. **a**, Conversion of holo T1 (grey) to loaded (red) in the presence of Cy1 WT. **b**, Concomitant response of Cy1 in presence of holo-T1 (blue) and loaded-T1 (pink). **c**, Conversion of holo-T1 (gold) to loaded-T1 (blue) in the presence of Cy1 D391N. **d**, Concomitant response of D391N in presence of holo-T1 (dark grey) and loaded-T1 (dark red). Bar charts in **a** and **c** show the percentage of substrate-loaded T1 when reaching a steady state, averaged over five T1 residues. Error bars indicate the standard deviation to the mean (see SI Methods). The signals of the minor conformer detected for wild-type Cy1 were not detected for D391N, which displayed only a few new signals that were too weak to be analyzed.

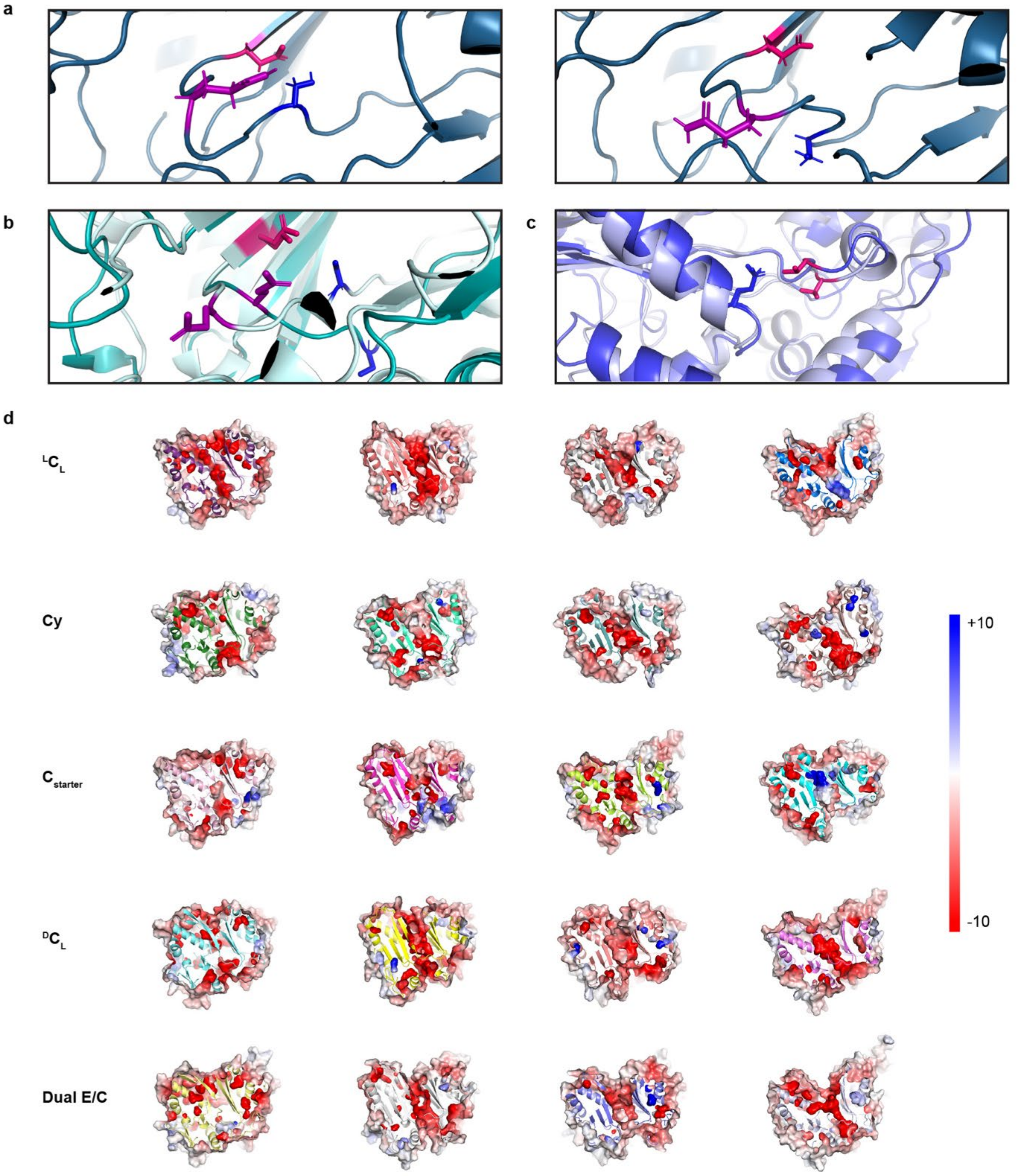

**SI Figure 10. Transient sidechain interactions and modulation of electrostatic potential in fluctuating conformations**. **a**, D391 (pink) transiently interacts with S383 (blue) in the Cy1 NMR structural ensemble as the conformation of Q387 (purple) changes dramatically. **b**, Similar structural fluctuations occur in the crystallographic ensemble of C-domain structures. A consensus sequence (SI Methods) for cyclization domains was used. **c**, A second example at the donor site, highlighted with a consensus sequence for condensation domains (position 287 in Cy1). Distinct conformations (light and dark blue) provide transient interactions between T272 (hot pink) and Q317 (blue). In **a**, **b**, and **c**, pink or purple highlight residues that impacted function when mutated, and blue highlight transient partners. The conformations correspond to pdbs 6p1j and 5t3e in **b**, and 4jn3 and 6p1j in **c**. **d**, Electrostatic potentials with cross-sections cut right above the β-strand that features D391 (in Cy1) in different consensus sequences (rows) and different conformations (columns). $^{L}C_{L}$: condensation domains with L-amino acids at both donor and acceptor sites; Cyc: cyclization domains; $C_{starter}$ starter C domain in NRPS modules; $^{D}C_{L}$: with donor accepting D-amino acids; Dual E/C: C domains with additional epimerization function. Columns report on conformations provided by 4jn3[5], 1l5a[13], 6p1j[14], and 5t3d[15], from left to right, which were also used in Fig. 3. Red: negative. Blue: positive. The scale on the right has units of kT/e. All cross-sections are at the same cutting plane.

#### SI DISCUSSION: Implications for Nonribosomal Peptide Synthetase mechanisms

Although our aim was not to determine molecular mechanisms in a traditional manner, and our main text focuses on the function of dynamics in a rather general manner, our results provide important insights on NRPS molecular mechanisms; notably regarding the remodelling of dynamic NRPS landscapes or because they explain otherwise confounding results in previous studies of the C-domain family.

Ever since the determination of the first structure of an NRPS module by Marahiel and co-workers,[16] it became clear that NRPS domains do not adopt a rigid quaternary architecture. Notably, T domains would need to visit catalytic domain partners in a series of sequential interactions, as further supported by NMR studies that demonstrated competing domain interactions involving T domains.[17] More recently, snapshots of modules capturing different stages of NRPS synthesis confirmed this hypothesis, notably through the work of Gulick and co-workers[15,18] and Schmeing and co-workers.[3,19,20] To achieve each synthetic step, NRPSs would then either need to transition between sequential, stable conformations or remain flexible throughout synthesis. Negative stain electron microscopy (EM) and cryo-EM data indicated that the second proposition is likely true as various modules or multi-modules displayed heterogeneous domain orientations[15,21]. The next question is then whether domains always randomly visit each other or synthesis remodels the NRPS dynamic landscape to promote interactions necessary for the next synthetic step. In the first model, a holo-T domain could visit a C or Cy domain, in which case no reaction would occur simply because no tethered building blocks are present, and the assembly line would still be functional. In contrast, in the second model, domains may interact randomly and fleetingly with each other but engage only for productive interactions. Here, a holo-T domain should only engage with an adenylation domain, and T-domains would only engage with C or Cy domains when they harbour the proper cargo. The molecular basis for cargo recognition should then be determined so that it may be preserved in engineered NRPSs, where exogenous substrates are introduced.

The chemical modifications of T domains imparted by NRPS synthesis appear to remodel the NRPS dynamic landscape to favour interactions with the domain catalysing the next modification. Thus, apo T domains bind tightly to phosphopantetheinyl transferases[22] but weakly to adenylation domains,[23] which gain affinity for T domains upon phosphopantetheinylation[23] and upon addition of aminoacyl adenylate mimics.[24,25] Accordingly, we found that Cy1 only engages with its partner donor T domain, T1, when T1 harbours a substrate, thus preventing unproductive interactions with a holo domain. The results are in agreement with those of Aldrich and Co., who highlighted that substrates were needed on the donor EntB to probe binding between EntB and EntF.[26] This observation is particularly relevant to engineering NRPS domain communication through domain swaps. Here, domains or groups of domains belonging to different systems are swapped and point-site mutations are introduced to restore communication between exogenous domains. Engineered systems must then not only enhance the affinity between exogenous domains, but ensure proper tuning of this affinity during synthesis, i.e as T-domains get modified. Engineering high affinity between exogenous domains may stall the NRPS machinery in the corresponding conformation. Determining how substrates, or their absence, tune domain/domain affinities is hence paramount to rational NRPS engineering, and our results provide some preliminary insights on this molecular cross-talk.

The salicylate tethered to T1 must interact with the surface of Cy1 and pass through a dynamic gate. The centre of the tunnel has been the focus of finding determinants for substrate recognition as it harbours the active site for peptide bond formation and cyclodehydration.[8,9,27] Although this region is necessarily critical for catalysis, our results indicate that it is not the only region of importance for substrate probing. We found that Cy1 binds only to salicylate-loaded T1 (Fig. 3f) but not to holo-T1, demonstrating that the substrate must interact with Cy1. We do not see any spectroscopic changes when Cy1 is presented to holo-T1 (SI Fig. 6a). NMR signals report on binding events described by dissociation constants, $K_D$s, ranging from nM to mM. Complexes with $K_D$s in the nM range lead to new signals, those in the μM range lead to reductions in signal intensities, and those in the mM range lead to changes in NMR signal frequencies. The emergence of new Cy1 signals in the presence of loaded-T1 is indicative of a stable complex. Contrastingly, if the phosphopantetheine arm was probed within Cy1's tunnel, we should see perturbations of Cy1 NMR signals when it is presented to holo-T1, presumably affecting residues within the tunnel, but also those at the T domain donor binding site as they would then interact with T1. That we see no perturbations in Cy1 while probing with holo-T1, indicates that Cy1 recognizes the absence of substrate at the end of the phosphopantetheine arm through a molecular encounter with an affinity reflecting a $K_D$ larger than mM, which could reasonably only occur at the surface of Cy1. This finding is complementary to and substantiated by recent observations made by Cryle and Co. who observed that the phosphopantetheine arm of a holo-T domain interacted with surface residues of the acceptor site of a condensation domain and only penetrated the domain's tunnel in the presence of a substrate.[7] This observation is consistent with a model wherein the T domain phosphopantetheine arm is probed at the surface of C domains and only enters the tunnel when it harbours a substrate. As mentioned in the main text, Aldrich and co-workers detected binding between EntB and EntF, which interact as a T-domain/C-domain pair, only when EntB was loaded with a substrate, again in agreement with our findings.[26] We do emphasize that our discussion does not allude to determinants of substrate specificity but to the discrimination between holo and loaded T-domains, although it stands to reason that, while substrate specificity may be governed within the buried active site, an interaction at the surface of the domain that precedes substrate funnelling is likely to also play a role in substrate specificity. We did not seek to determine the site for substrate recognition as we focused on demonstrating the role of global dynamics in governing this recognition. However, the structural fluctuations we observe at the donor site (SI Fig. 4a) identify regions that could serve as a dynamic gate. In Cy1, these regions are defined by loops L16 and L20, which represent the second and third crossing between the C-terminal and N-terminal regions and thereby delimit the floor and roof entrances to the donor site (in pink and purple, respectively, in Fig. 2 and SI Figs. 3 and 4). It will be critical to determine whether and which residues in these regions recognize specific substrates for a better understanding of both the gating mechanism and maybe of substrate specificity in Cy1.

Dynamics *within* cyclization or condensation domains may help remodel the dynamics *between* NRPS domains by promoting interactions with loaded T-domains and preventing interactions with holo T-domains. A key aspect of our results is to link molecular recognition and allosteric responses with Cy1's dynamics. The site of the mutation D391N is out of reach of T1 or its prosthetic group (Fig. 4a) and yet hampers the affinity of Cy1 for loaded T1, as evidenced by weaker and much fewer signals exhibiting a response when D391N is presented with loaded-T1 (SI Fig 9d). We demonstrated that this mutation not only impeded Cy1 global dynamics (Fig 4c, SI Fig. 8d), but also stabilized its fold (Fig. 4b and SI Fig. 8a-c). This observation supports a model where structural fluctuations within Cy1 are needed to efficiently discriminate between holo and loaded T1, such that substrate attachment tunes the affinity between both forms. While our focus necessarily required an approach using domains *in trans*, NRPS domains including those in our model system often interact *in cis*. In the context of multi-domain assembly lines, binding events correspond to interactions between domains that are also reflected by changes in orientations between domains. As mentioned above, NRPS modules display heterogeneous domain orientations and, hence, a variety of domain interactions. Cy1's internal structural dynamics would then allow for remodelling the NRPS dynamic landscape such that only loaded-T1 would engage with Cy1 while holo-T1 is left accessible to other domains, i.e. YbtE. Although challenging, it will be important to account for C or Cy-domain internal dynamics in future studies of multi-domain assemblies. Notably, although we are educated to only consider surface residues when studying interactions between domains, the D391N mutation demonstrates that residues within C-domain tunnels could affect product formation not only through direct contacts with substrates but also by affecting interactions between domains.

Cyclization domains appear to possess a secondary function for substrate editing, like C domains do. Recently, Cryle and Co-workers have demonstrated that condensation domains could hydrolyse the thioester bonds linking substrates to their dedicated thiolation domains.[28] This function provides a kinetic constraint on NRPS assembly lines as both acceptor and donor substrates must reach C domain active sites before their substrates are hydrolysed. In previous work, we loaded T1 up to near completion with various concentrations of YbtE, which only altered the time needed to reach (near) complete loading. However, in presence of Cy1, the population of loaded-T1 appears limited by the balance between the concentrations of YbtE and Cy1 and, hence, Cy1 appears to accelerate the hydrolysis of salicylate tethered to T1. Accordingly, in our first attempt to load T1 *in situ* in presence of Cy1, we could only reach a steady conversion of 58% when 560 μM T1 was in presence of 500 μM Cy1 with 100 nM YbtE. We put this observation to our advantage as we varied the concentration of YbtE to obtain a correlation between the populations of loaded-T1 and the minor conformer that we detected for Cy1 (Fig. 3e). Here, Cy1 was present at only 100 μM, and 80% of conversion could be reached with 200 nM YbtE. Future studies will be needed to verify and quantify cyclization domain hydrolase activity and take it into account in the context of NRPS assembly lines.

The allosteric response of Cy1 through substrate recognition resolves puzzling observations that point at T domain communication across C-domains. Substrate-loaded Co-enzyme A (CoA) and derivatives thereof are commonly used as surrogates of T-domains in mechanistic studies of NPRSs or related fatty acid or polyketide synthases. Schmeing and co-workers noted that C-domains will only accept surrogates when the second substrate is tethered to a T-domain,[27] an observation difficult to explain with rigid structures featuring open tunnels. Indeed, a T-domain sits 40 Å apart from the remaining opening where the CoA surrogate enters, and hence the T-domain cannot impact the surrogate's affinity for C-domains through a direct structural interaction. The allosteric response we observed in Cy1 (Fig. 3g) provides a solution to this conundrum: a stable interaction with a loaded T-domain provides a molecular response that reaches the remote partner T-domain site. Apparently, this response can facilitate the access of phosphopantetheine arms not attached to T-domains as the other site is occupied by a T-domain. It will be critical to establish whether this communication translates into a coupling of substrate or domain specificity for successful engineering of NRPSs; C-domains may need productive interactions with both their donor and acceptor T-domains and tethered substrates for function.

The global dynamics we observed for Cy1 may be used to funnel loaded phosphopantetheine arms into its tunnel. Although the open tunnels that were observed across many C- and Cy-domains can often accommodate phosphopantetheines with substrates, how these moieties find their way into the tunnel and how extended products leave the tunnel remains unsolved. Indeed, T-domains do not present their substrates through rigid extended arms. We previously demonstrated that T1 samples an equilibrium between a docked form and an undocked form with a disengaged, disordered phosphopantetheine arm.[12] Whichever form Cy1 encounters, the phosphopantetheine arm must be guided into the active site to reach the conformations captured by X-ray crystallography. Taking into consideration a dynamic gate at the donor site paired with a global malleability, we suggest a mechanism wherein the substrate is probed at the surface near the dynamic gate and coupled, global dynamics may then be used to help funnel the substrate into the active site by modulating the shape of the tunnel. This hypothesis is supported by a remarkable study performed on a related domain in enacyloxin synthesis.[29] This domain belongs to the C-family but only needs to interact with a donor T-domain as it condenses its cargo with a free ligand to release the product. Remarkably, and perhaps accordingly, the domain was found with an open latch, i.e. with a tunnel replaced by a channel or cavity defined by the N- and C-terminal regions, which are now disengaged. Accelerated molecular dynamics not only revealed closing of the latch but also concomitant opening of an occluded acceptor site paired with closing of the donor site, in a mechanism reminiscent of the gating mechanism we established experimentally paired with the formation of an occluded tunnel. Global C-domain dynamics may similarly lead to tunnel dynamics that could help thread phosphopantetheines and substrates towards the active site. Indeed, C-domain structures often exhibit different conformations of the donor site as well as varying degrees of tunnel openness and constrictions. In Cy1, the snapshots provided by tunnels calculated with our NMR ensemble (Fig. 2 and SI Fig. 2c), the dynamic footprint of Cy1, which depicts a malleable tunnel (Fig. 3), and a response to loaded-T1 that encompasses the entire tunnel together support an active role of Cy1 dynamics in promoting the productive complex between Cy1 and loaded-T1.

The opening of the latch in C-domains is itself a recurrent point of discussion in the community because it could explain how intermediates leave C-domains following extension. Without latch opening, an extended intermediate would have to thread through the tunnel as the acceptor T domain they are tethered to disengages the C-domain to visit its next partner. Considering the variety of products and intermediates that can be found in NRPS systems, this model seems unlikely. Latch opening has been mentioned for a long time, as low density was observed in that region,[30] but a fully disengaged latch had not been observed until the work of

Kosol et al.[29] However, this domain only interacts with a single T-domain and cannot be used as a paradigm for C or Cy domains. In solution, Cy1's latch is poorly defined and subject to structural fluctuations that may reflect transient unpairing. However, relaxation dispersion does not point at a stable minor open state since the changes in chemical shifts captured by relaxation dispersion with a two-state model in that region (residues 368 and 375) are too small to reflect disengagement. That is, Cy1 does not provide a lowly populated open state for release of the extended product. Instead, formation of the extended product may disrupt the aforementioned transient interactions at the latch to promote opening of Cy1 before the acceptor T-domain and its extended cargo disengage.

Finally, the global dynamics we determined for Cy1 in solution explain in part confounding responses towards single-point mutations in condensation and cyclization domains. Both domains have been extensively probed through mutagenesis and some conserved positions appear to only be interpretable for specific systems.[8,9,27] Traditionally, the impact of a mutation is interpreted against a static backdrop through local structural considerations such as the interaction between a sidechain and substrates or the interaction between sidechains that maintain structural integrity. Because different condensation domains have crystalized in distinct conformations, the impact of mutating some sites may appear to only be interpretable for a given system, where the conformation highlights a given interaction. However, our results indicate that the changes in conformations captured by crystallography reflect structural fluctuations in solution.

In a global dynamic environment, a sidechain may experience different molecular contacts in different conformations sampled during structural fluctuations. A mutation could either prevent access to certain conformations, or lock the protein in a stable conformation, or enhance structural fluctuations. Notably, we observed that, although D391 has a sidechain available for substrate interactions in the structures of EpoB and BmdB cyclization domains (D449 and D1226, respectively), in the NMR ensemble it is sometimes found poised to interact with the conserved serine S383 (SI Fig. 10a). The global response we have observed on D391N may well be in part due to disrupting transient interactions with S383, although future studies will be needed to demonstrate this hypothesis. To illustrate these effects in a more general manner, we threaded C-domain and Cy-domain consensus sequences (SI Methods) onto conformations seen by crystallography (SI Figs. 10b, c) and found that the crystallographic ensemble replicates the findings made with the NMR ensemble. This observation strengthens both that the changes of conformations captured by crystallography reflect structural fluctuations in solution, and that the dynamics we observe for Cy1 are common in the C-domain family. Thus, the aspartate/serine pair is again seen to interact transiently. Further, another conserved residue, Q387 in Cy1 (purple), undergoes similar conformational changes in the crystallographic (SI Figs. 10b) and NMR ensembles (SI Figs. 10a), moving in and out of the aspartate/serine interaction pair. We provide a second example, this time at the donor site and using a C-domain consensus sequence. T272 follows a conserved arginine and, although T272 is itself not conserved (it is Q287 in Cy1), mutation drastically affected condensation in VibH (T272 in consensus is W264 in VibH).[13] This region is shown to be dynamic by relaxation dispersion, which is reflected by alternative conformations in the crystallographic ensemble (dark and light blue in SI Figure 10c). Thus, T272 (in pink in SI Figure 10c) is seen to transiently interact with Q317 (in blue in SI Figure 10c), and mutagenesis will disrupt this transient interaction.

Mutations at the positions highlighted may impart responses similar to that we observed for D391. The altered side chains may no longer be able to transiently interact with partner side chains and global dynamics become disrupted. We by no means suggest that the only impact of mutating these residues is to disrupt dynamics. In fact, in many conformations of Cy1's NMR ensemble, the carboxylic group of D391 is indeed available to interact with an intermediate, as proposed by two groups including that of a co-author.[8,9] Similarly, Q387 offers its sidechain to the donor binding site in some conformations, and mutagenesis may disrupt intermolecular contacts. However, our results demonstrate that the molecular interpretation of a point-site mutation should account for dynamics, which may explain challenges in interpreting past results prior to this observation. Critically, while sites of interests are identified through inspections of local features within domains, function is traditionally assayed through multi-domains or modules, and our findings demonstrate that a site within the core of the protein and remote from binding sites may nevertheless impact domain communication. Indeed, here, we strived to provide a readout that isolates a specific response within Cy1 through our *in situ* NMR experiments rather than relying on product formation as our observations rapidly indicated that interpretation of the latter would be challenging. Thus, it will be important to determine whether and how communication with domains other than T-domains may be impacted by C-domain dynamics and by mutations that disrupt them.

In the course of analysing consensus C and Cy sequences, we discovered a site displaying a negative electrostatic potential that seems to elongate and contract in a conformation dependent manner. In its extended form, the region stretches from the beginning of $\alpha4$, which interacts with the donor T domain, to helices $\alpha1$ and $\alpha10$, which define the binding site for the acceptor T domain. The residues that define this region are not exclusively found in the tunnel in that conformation. We observe that the negative electrostatic potential is relieved as the conformation changes (SI Fig. 10d, first row, left to right). Thus, the global dynamics we observe are accompanied by a modulation of this negative electrostatic potential, which in turn may be tuned by interactions with partner substrates and domains. That is, the modulation of the electrostatic potential in that region may provide a means to tune the dynamic mediated allosteric communication we have observed. Interestingly, although we also observed a similar occurrence for other members of the C-domain family (SI Fig. 10d, second to fifth rows, left to right), such as C starter domains, ${}^{D}C_{L}$, and dual epimerization/condensation domains, their fine features differ markedly when they are displayed on the same common conformation (SI Fig. 10d columns). It will be interesting to probe how these variations may reflect on the diverse functions of these domains.

Overall, our results bring new perspectives to interpret NRPS molecular mechanisms in light of the allosteric function of C-domain dynamics and may open doorways towards successful approaches to engineer exogenous substrate recognition into C or Cy domains.

#### SI METHODS

##### ***Expression and purification of Ybt Cy1 with various labeling schemes.***

The cloning, expression, and purification of Ybt Cy1, a 52 kDa cyclization domain excised from the yersiniabactin synthetase NRPS HMWP2, has been described previously.[31,32] The purification scheme is described here in brief, with minor modifications compared to the previously reported protocols. BL21 (DE3) cells (Novagen) transformed with the pET30a-Cy1H6 plasmid were grown in various minimal media depending on the desired isotopic labeling scheme. Cells were harvested by centrifugation and lysed using a microfluidizer, followed by centrifugation to remove cellular debris. The soluble fraction was filtered and subjected to affinity purification using a HisTrap HP column (GE Healthcare, Sweden) to capture and elute Cy1 with its hexahistidine (H6) C-terminal tag. The eluted fractions were confirmed by SDS-PAGE analysis and dialyzed overnight, followed by concentration and purification using size-exclusion chromatography (SEC) in SEC buffer (20 mM sodium phosphate buffer, 100 mM NaCl, 1 mM EDTA, 5 mM DTT, pH 7).

The following protocol was implemented to facilitate exchange of Cy1 solvent-protected amides from deuterons to protons. SEC-purified Cy1 was concentrated and allowed to back exchange deuterons for protons in degassed NMR exchange buffer (50 mM Tris pH 8.5 at room temperature, 10 mM NaCl, 1 mM EDTA, 10 mM DTT) for approximately 5 days at room temperature in a sealed container. Cy1 was then concentrated using an Amicon Ultra centrifugal filter (10K NMWL, Millipore, Ireland), and subjected to a final size exclusion chromatography step in SEC buffer. The eluted protein was then concentrated and buffer exchanged into the appropriate NMR buffer by diafiltration using a 10K molecular weight cut-off centrifugal filter (2000 x g at 18 °C).

The following isotopic labeling schemes were used in this study: CDN (uniform $^{13}C$, $^{2}H$, $^{15}N$), CDN-ILV ($^{13}C$, $^{2}H$, $^{15}N$ for backbone; $^{13}C$, $^{1}H$ for isoleucine δ1, leucine δ1 and δ2, and valine γ1 and γ2 methyl groups only), 70DCN ($^{13}C$, 70% $^{2}H$ / 30% $^{1}H$, $^{15}N$ backbone and sidechains), DN-ILV stereo ($^{12}C$, $^{2}H$, $^{15}N$ for backbone and sidechains, except $^{13}C$, $^{1}H$ for isoleucine δ1, leucine δ2, and valine γ2 methyl groups), DN-FYILV ($^{12}C$, $^{2}H$, $^{15}N$ for backbone and sidechains, except $^{12}C$, $^{1}H$, $^{15}N$ for phenylalanine and tyrosine, and $^{13}C$, $^{1}H$ for isoleucine δ1, leucine δ1 and δ2, and valine γ1 and γ2 methyl groups).

The procedure and materials used for samples with CDN and CDN-ILV isotopic labeling have been published previously[32-34]. The recipe for growth media for the DN-ILV stereo labeling is similar to that of DN-FYILV samples described in [32] without any $^{15}N$-phenylalanine or $^{15}N$-tyrosine added to the growth media. Additionally, the precursors for stereospecific isotopic labeling were 2-($^{13}C$)-methyl-4-($^{2}H_3$)-acetolactate (leucine and valine) and $^{13}C$-methyl α-ketobutyric acid (isoleucine).[35] Isotope precursors were added to the expression media prior to induction, when the optical density at 600 nm ($OD_{600}$) reached ~ 0.5. The acetolactate precursor was produced by incubating ethyl-2-hydroxy-2-($^{13}C$)-methyl-3-oxobutanoate in 10 ml $D_2O$ with a pD of ~ 13 for an hour and neutralized before use. The 70DCN labeling scheme used the same components as for the CDN sample, except the M9 minimal media was composed of 70 % (v/v) $D_2O$ and 30 % (v/v) $H_2O$.

***Cloning of Ybt Cy1 D391N mutant.***

All Cy1 mutant data were recorded on a single-point Asp to Asn mutation in Cy1 WT at position 391 (position 490 in the full-length Ybt HMWP2 numbering). Mutagenesis was performed using PCR guided site-directed mutagenesis utilizing the wild-type Cy1 construct[32] and primers designed as described in Liu et al.[36] i.e. 5'-gtctggataaatcatctggcgttcgagcatcacggcgaggtc-3' (forward primer) and 5'-cagatgatttatccagacctgcggcgtttgcgagatgccccattc-3' (reverse primer). The final purified template is identical to the WT Cy1 construct with the single-point D391N mutation.

***Expression and purification of Cy1 D391N Mutant.***

The Ybt Cy1 D391N mutant used in NMR experiments was uniformly isotopically labeled with $^{2}H$, $^{15}N$, and $^{13}C$ (CDN) and expressed and purified with a protocol similar to WT. For thermal denaturation studies, unlabeled Cy1 (WT and D391N mutant) were expressed in LB media containing kanamycin (50 mg/L). Based on optical measurements, the purification yielded 50 mg of CDN-labeled protein. The pooled fractions were concentrated to 35 mL in preparation for $^2H$ to $^1H$ back exchange. The protein was back exchanged into 100 mL of exchange buffer (50 mM Tris, 10 mM NaCl, 10 mM EDTA, 10 mM DTT, pH 8.5) at room temperature for approximately 9 days. Following back-exchange, the protein was concentrated to 3 mL and purified by size-exclusion chromatography using 1 mL injection volumes.

Preparation of the final NMR sample for the Cy1 D391N construct proceeded by buffer exchanging the final purified protein into NMR titration buffer (50 mM ACES, 10 mM NaCl, 2 mM $MgCl_2$, pH 7.0 at room temperature). The protein was finally concentrated to approximately 400 μL corresponding to a Cy1 D391N concentration of ~ 900 μM. The final NMR sample was made in a Shigemi NMR tube (Sigma/Aldrich) with a Cy1 D391N concentration of 796 μM containing 5% $D_2O$, 250 μM DSS, layered with argon gas, and sealed with parafilm.

Purification of unlabeled Cy1 (WT and D391N mutant) followed a similar protocol, but with the following adaptations. Lysis of cells was performed using an ultrasonic liquid processor (Misonix). The lysate was kept on ice at all times during sonication. The purification then followed that of the mutant above with no need for a back-exchange step. From a 1-L growth, 169 mg of Cy1 WT and 128 mg of Cy1 D391N. The proteins were stored dilute in size-exclusion buffer before being flash-frozen and kept at -80 °C. Before thermal denaturation studies, the proteins were thawed at room temperature and buffer exchanged into NMR buffer (50 mM ACES, 10 mM NaCl, 2 mM $MgCl_2$, pH 7.0 at room temperature).

***NMR Experiments and Acquisition Parameters.***

We provide below tables of 3D and 4D experiments recorded with various samples at various field strengths. Acquisition parameters are listed when not mentioned in previous published studies. Samples are defined according to their labeling schemes. CDN: U-$^{13}$C -$^{15}$N-$^{2}$H; 70DCN: U-$^{13}$C -$^{15}$N-70%$^{2}$H; CDN-ILV: $^{1}$H-$^{13}$C-Me- δ1I,L,V-U-$^{13}$C-$^{2}$H-$^{15}$N; DN-FYILV: $^{1}$H-$^{13}$C-Me- δ1I,L,V-U-$^{2}$H-$^{15}$N; DN-ILV stereo: $^{1}$H-$^{13}$C-Me-δ1I,δ2L,γ2V-U-$^{2}$H-$^{15}$N. Sample concentrations used were as follows: for wild type Cy1 CDN (588 μM), Cy1 CDN (RDC) (429 μM), Cy1 CDN (Hahn-Echo) (600 μM), 70DCN (986 μM), CDN-ILV (586 μM), DN-FYILV (350 μM), DN-ILV stereo (600 μM). For Cy1 D391N, CDN (796 μM) was used in backbone assignments and CDN (122 μM) was used in dynamics experiments. TROSY versions of NMR pulse sequences were used. Non-uniform sampling (NUS) sampling factor is noted when it was used.

***Table S2. NMR Acquisition Parameters: Backbone Assignment***

| Labeling Scheme | Experiment & Field Strength (MHz) | NS / Recycling Delay (s) | TD | Spectral Width (ppm) @ Carrier Frequencies (ppm) |
|---|---|---|---|---|
| CDN | HNCA: 800 | 16/1 | 2080 ($^{1}$H detected) x 350 ($^{15}$N) x 200 ($^{13}$C), 2 % sampling | 20 @ 4.8 ($^{1}$H), 31 @ 58 ($^{13}$C), 35 @ 118 ($^{15}$N) |
| CDN | HNCO: 800 | 8/1 | 2080 ($^{1}$H detected) x 400 ($^{15}$N) x 100 ($^{13}$C), 3.125 % sampling | 20 @ 4.8 ($^{1}$H), 16 @ 176 ($^{13}$C), 35 @ 118 ($^{15}$N) |
| CDN | HN(CA)CO: 800 | Uniform sampling Ref.[33] | | |
| CDN | HN(CO)CA: 800 | Uniform sampling Ref.[33] | | |
| CDN | HN(CA)CB: 800 | 16/1 | 1650 ($^{1}$H detected) x 110 ($^{15}$N) x 148 ($^{13}$C) | 18.75 @ 4.8 ($^{1}$H), 62 @ 44 ($^{13}$C), 35 @ 118 ($^{15}$N) |
| CDN-ILV | HN(COCA)CB: 600 | 192/1 | 2048 ($^{1}$H detected) x 45 ($^{15}$N) x 75 ($^{13}$C), 15 % sampling | 18.75 @ 4.8 ($^{1}$H), 62 @ 44 ($^{13}$C), 35 @ 118 ($^{15}$N) |

| | | | | |
|---|---|---|---|---|
| CDN-ILV | Ala-HN(CA)CB: 600 | 112/1 | 2048 ($^1$H detected) x 45 ($^{15}$N) x 25 ($^{13}$C), 10 % sampling | 16@ 4.8 ($^1$H), 15 @ 18.5 ($^{13}$C), 33 @ 118 ($^{15}$N) |
| CDN-ILV | Gly-HNCA: 600 | 128/1 | TD 2048 ($^1$H detected) x 45 ($^{15}$N) x 25 ($^{13}$C), 10 % sampling | 16 @ 4.8 ($^1$H), 20 @ 45 ($^{13}$C), 33@ 119 ($^{15}$N) |
| CDN-ILV | Ser/Thr-HN(CA)CB: 600 | 224/1 | 2048 ($^1$H detected) x 45 ($^{15}$N) x 25 ($^{13}$C), 10 % sampling | 16@ 4.8 ($^1$H), 20 @ 18.5 ($^{13}$C), 33 @ 118 ($^{15}$N) |
| CDN-ILV | Ser/Thr-HN(COCA)CB: 600 | 112/1 | 2048 ($^1$H detected) x 45 ($^{15}$N) x 25 ($^{13}$C), 10 % sampling | 16@ 4.8 ($^1$H), 20 @ 18.5 ($^{13}$C), 33 @ 118 ($^{15}$N) |
| 70DCN | HACACO: 600 | 64/1 | 1024 ($^{13}$C detected) x 48 ($^1$H) x 84 ($^{13}$C) | 40 @ 176 ($^{13}C_{direct}$), 4.5 @ 4 ($^1$H), 35 @ 55.5 ($^{13}$C) |
| 70DCN: | HNCO: 800 | 96/1 | 1600 ($^1$H detected) x 60 ($^{15}$N) x 45 ($^{13}$C), 5 % sampling | 20 @ 4.8 ($^1$H), 15 @ 175.5 ($^{13}$C), 32 @ 118 ($^{15}$N) |
| 70DCN | HNCA: 800 | 256/1 | 1600 ($^1$H detected) x 60 ($^{15}$N) x 52 ($^{13}$C), 5 % sampling | 20 @ 4.8 ($^1$H), 28 @ 55.5 ($^{13}$C), 32 @ 118 ($^{15}$N) |
| 70DCN | HN(CA)CO: 800 | 96/1 | 1600 ($^1$H detected) x 60 ($^{15}$N) x 45 ($^{13}$C), 5 % sampling | 20 @ 4.8 ($^1$H), 15 @ 175.5 ($^{13}$C), 32 @ 118 ($^{15}$N) |
| 70DCN | HN(CO)CA: 800 | 256/1 | 1600 ($^1$H detected) x 60 ($^{15}$N) x 52 ($^{13}$C), 5 % sampling | 20 @ 4.8 ($^1$H), 28 @ 55.5 ($^{13}$C), 32 @ 118 ($^{15}$N) |
| CDN (D391N) | 2D HN-TROSY | 16/1 | 2048 ($^1$H detected) x 300 ($^{15}$N) | 16 @ 4.7($^1$H), 31 @ 118 ($^{15}$N) |
| CDN (D391N) | HNCO 600 MHz | 16/1 | 2048 ($^1$H detected) x 80 ($^{15}$N) x 60 ($^{13}$C) | 16 @4.7 ($^1$H), 31 @ 118 ($^{15}$N), and 16 @176 ($^{13}$C) |
| CDN (D391N) | HNCA 600 MHz | 16/1.05 | 2048 ($^1$H detected) x 90 ($^{15}$N) x 100 ($^{13}$C) | 16.02 @4.7 ($^1$H), 31 @ 118($^{15}$N), and 28 @ 56($^{13}$C) |

#### *Table S3. NMR Acquisition Parameters: Signal Assignment (Methyl Sidechain)*

| Labeling Scheme | Experiment & Field Strength (MHz) | NS / Recycling Delay (s) | TD | Spectral Width (ppm) @ Carrier Frequencies (ppm) |
|---|---|---|---|---|
| CDN-ILV | HMCMCBCA: 800 | Ref.[31] | | |
| CDN-ILV | HMCMCGCBCA: 800 | Ref.[31] | | |
| CDN-ILV | HCCH TOCSY: 800 | Ref.[31] | | |
| CDN-ILV | HC(CCO)NH: 600 | Ref.[34] | | |

***Table S4. NMR Acquisition Parameters: Structure Calculation (Distance Restraints) & Methyl Sidechain (ILV) Signal Assignment***

| Labeling Scheme | Experiment & Field Strength (MHz) | NS / Recycling Delay (s) | TD | Spectral Width (ppm) @ Carrier Frequencies (ppm) |
|---|---|---|---|---|
| DN-FYILV | 4D time-shared (TS) $^{15}$N/$^{13}$C HSQC-NOESY-TROSY/ HSQC: 800[a] | 8/1 | 1620 ($^{1}$H detected) x 110 ($^{15}$N / $^{13}$C TROSY/HSQC) x 100 ($^{1}H_{NOESY}$) x 30 ($^{15}$N / $^{13}$C HSQC/HSQC), 1 % sampling | 16.875 @ 4.76 ($^{1}$H direct), 21 @ 18.5 ($^{13}$C), 35@ 119 ($^{15}$N), 4.75 @ 8 ($^{1}$H) |
| DN-FYILV | TS-TROSY/HSQC NOESY: 600 | Ref.[32] | | |
| DN-FYILV | TS-NOESY - HSQC/TROSY: 600 | Ref.[32] | | |
| DN-ILVstereo | TS-TROSY/HSQC NOESY: 950[b] | 32/1 | 1024 ($^{1}H_{noe}$ detected) x 80 ($^{15}$N /$^{13}$C) x 200 ($^{13}$C) | 16 @ 4.7 ($^{1}H_{noe}$ direct), 32 @ 118 ($^{15}$N time shared) and 23 @ 18 ($^{13}$C time shared), 6.5 @ 4.7 ($^{1}$H) |
| CDN | $^{15}$N TROSY-NOESY: 800[c] | 8/1 | 1650 ($^{1}$H detected) x 180 ($^{15}$N) x 124 ($^{13}$C) | 18.75 @ 4.75 ($^{1}$H direct), 13 @ 4.75 ($^{1}H_{noe}$) 35 @ 118 ($^{15}$N) |
| CDN (RDC) | HNCO:600 Interleaved TROSY and non-TROSY versions (see details in later section) | 64/1 | 2048 ($^{1}$H detected) x 65 ($^{15}$N) x 35 ($^{13}$C), 30 % sampling | 16 @ 4.8 ($^{1}$H), 16 @ 176 ($^{13}$C), 33 @ 118.5 ($^{15}$N) |

[a] NOE mixing time: 150 ms

[b] NOE mixing time: 40 and 150 ms (two sets)

[c] NOE mixing time: 100 ms

***Table S5. NMR Acquisition Parameters: Dynamics***

| Labeling Scheme | Experiment & Field Strength (MHz) | NS / Recycling Delay (s) | TD | Spectral Width (ppm) @ Carrier Frequencies (ppm) |
|---|---|---|---|---|
| DN-ILVstereo | $^{15}$N RD-CPMG 950[a] | 32/3.5 | 2048 ($^{1}$H detected) x 260 ($^{15}$N) | 18 @4.75 ($^{1}$H detected) x 30.5 @ 117.75 ($^{15}$N)<br><br>13 points (34.92 – 961.54 Hz), CT = 28.8 ms |
| DN-ILVstereo | $^{15}$N RD-CPMG, 600[b] | 32/3.5 | 2048 ($^{1}$H detected) x 300 ($^{15}$N) | 18 @4.75 ($^{1}$H detected) x 30.5 @ 117.75 ($^{15}$N) |

| | | | | |
|---|---|---|---|---|
| | | | | 15 points (29.07 – 1008.06 Hz), CT = 34.56 ms |
| CDN | Hahn Echo, 600[c] | 64/1 | 2048 ($^1$H detected) x 300 ($^{15}$N) | 16 @ 4.7 ($^1$H) 31 @ 118 ($^{15}$N) ppm, $^{13}$C decoupling @ 54 ppm |
| CDN (D391N) | Hahn Echo 600[d] | 256/1 | 2048 ($^1$H detected) x 300 ($^{15}$N) | 16 @ 4.7 ($^1$H), 31 @118 ($^{15}$N) ppm, $^{13}$C decoupling @ 54 ppm |

[a] 13 points with CPMG frequencies in the range 34.92 – 961.54 Hz and the reference (0 Hz), all collected in an interleaved manner. CT: Constant Time period.

[b] 15 points with CPMG frequencies in the range 29.07 – 1008.06 Hz and the reference (0 Hz), all collected in an interleaved manner.

[c] the delay $\tau$ = 10.8 ms[37]

[d] $\tau$ = 10.8 ms[37]

***Table S6. NMR Acquisition Parameters: Cy1 – T1 Complexes***

| Labeling Scheme | Experiment & Field Strength (MHz) | NS / Recycling Delay (s) | TD | Spectral Width (ppm) @ Carrier Frequencies (ppm) |
|---|---|---|---|---|
| Cy1 WT (CDN) free | HNCO: 600 | 16/3 | 2048 ($^1$H detected) x 90 ($^{15}$N) x 60 ($^{13}$C) | 16 @ 4.8 ($^1$H), 16 @ 176 ($^{13}$C), 31 @ 118 ($^{15}$N) |
| Cy1 WT (CDN) + T1 holo ($^{15}$N/$^1$H/$^{12}$C) | HNCO: 600 | 16/1 | 2048 ($^1$H detected) x 80 ($^{15}$N) x 60 ($^{13}$C) | 16 @ 4.8 ($^1$H), 16 @ 176 ($^{13}$C), 31 @ 118 ($^{15}$N) |
| Cy1 WT (CDN) + T1 holo or loaded ($^{15}$N/$^1$H/$^{12}$C) | 2D IDIS TROSY: 600 | 8/1 | 2048 ($^1$H detected) x 128 ($^{15}$N) | 16 @ 4.8 ($^1$H), 31 @ 118 ($^{15}$N), $^{13}$C decoupler at 176 ppm |
| Cy1 WT (CDN) + T1 loaded *in situ* ($^{15}$N/$^1$H/$^{12}$C) | HNCO: 600 | 16/3 | 2048 ($^1$H detected) x 90 ($^{15}$N) x 60 ($^{13}$C) | 16 @ 4.8 ($^1$H), 16 @ 176 ($^{13}$C), 31 @ 118 ($^{15}$N) |
| Cy1 WT (CDN) + T1 loaded ($^{15}$N/$^1$H/$^{12}$C), (before addition) | HNCO: 600 | 16/3 | 2048 ($^1$H detected) x 90 ($^{15}$N) x 60 ($^{13}$C) | 16 @ 4.8 ($^1$H), 16 @ 176 ($^{13}$C), 31 @ 118 ($^{15}$N) |

| | | | | |
|---|---|---|---|---|
| Cy1 WT (CDN) + loaded T1 before addition, ($^{15}N/^{1}H/^{12}C$), unloaded by SrfAD | HNCO: 600 | 16/3 | 2048 ($^{1}H$ detected) x 80 ($^{15}N$) x 60 ($^{13}C$) | 16 @ 4.8 ($^{1}H$), 16 @ 176 ($^{13}C$), 31 @ 118 ($^{15}N$) |
| Cy1 WT (CDN-ILV) + T1 loaded *in situ* ($^{15}N/^{2}H/^{12}C$) | TS-TROSY-HSQC: 600 | 16/1 | 2080 ($^{1}H$ detected) x 90 ($^{15}N$ / $^{13}C$ time shared) | 20 @ 4.8 ($^{1}H$), 21 @ 19 ($^{13}C$), 35 @ 118 ($^{15}N$) |
| Cy1 WT (CDN) + T1 loaded before addition, ($^{15}N/^{2}H/^{12}C$) | HNCO: 600 | 16/1 | 2048 ($^{1}H$ detected) x 80 ($^{15}N$) x 60 ($^{13}C$) | 16 @ 4.8 ($^{1}H$), 16 @ 176 ($^{13}C$), 31 @ 118 ($^{15}N$) |
| Cy1 WT (CDN) + T1 loaded before addition, ($^{15}N/^{2}H/^{12}C$) | HNCA: 600 | 32/1 | 2048 ($^{1}H$ detected) x 80 ($^{15}N$) x 80 ($^{13}C$) | 16 @ 4.8 ($^{1}H$), 28 @ 56 ($^{13}C$), 31 @ 118 ($^{15}N$) |
| CDN (D391N) + DN (T1): **For all D391N – T1 complex samples | 2D IDIS-TROSY: 600 | 16/1 | 2048 ($^{1}H$ detected) x 600 ($^{15}N$) | 16 @ 4.7 ($^{1}H$), 31@118 ($^{15}N$), $^{13}C$ decoupling at 176 ppm. |
| CDN (D391N) + DN (T1): Loaded T1 complex | HNCO: 600 | 32/1 | 2048 ($^{1}H$ detected) x 80 ($^{15}N$) x 60 ($^{13}C$) | 16 @ 4.7 ($^{1}H$), 31 @118 ($^{15}N$), and 16 @ 176 ($^{13}C$) ppm |
| CDN (D391N) + DN (T1): Loaded T1 complex | HNCA: 600 | 48/1 | 2048 ($^{1}H$ detected) x 80 ($^{15}N$) x 80 ($^{13}C$) | 16 @ 4.7 ($^{1}H$), 31 @ 118 ($^{15}N$), and 28 @ 56 ($^{13}C$) ppm |
| CDN (D391N) + DN (T1): Unloaded T1 complex after addition of SrfAD | HNCO: 600 | 40/1 | 2048 ($^{1}H$ detected) x 80 ($^{15}N$) x 60 ($^{13}C$) | 16 @ 4.7 ($^{1}H$), 31 @ 118 ($^{15}N$), and 16 @ 176 ($^{13}C$) ppm |

***NMR Data Processing and Analysis***

All NMR data was processed using NMRPipe [38] and analyzed in CARA [39]or SPARKY.[40] Covariance maps to facilitate backbone and side-chain assignments were calculated as published previously.[41,42] Non-uniform sampling schedules were made with PoissonGap software and the data were processed with istHMS.[43] Relaxation dispersion data was analyzed using ChemEx.[44] Ultimately 93% of Cy1 backbone resonances were assigned using combinations of various isotopically labeled samples and procedures. A structural model of Cy1 created using the Cy1 sequence and the crystal structure of the EpoB cyclization domain[9] was used along with NOESY 3D and 4D spectra for identifying distance-based backbone as well as side-chain assignments for phenylalanine, tyrosine, and methyl groups.

***NMR Measurements of Ybt Cy1 D391N (CDN).***

All D391N NMR measurements were carried out at 25 °C on a Bruker 600 MHz AVANCE III spectrometer equipped with a QCI cryoprobe™. The uniformly labeled $^{15}N$, $^{13}C$, $^{2}H$ (CDN) sample was used in NMR buffer.

*Backbone Assignment*: To assign the resonances of the mutant HNCO and HNCA triple-resonance experiments were collected (see section 'NMR Experiments and Acquisition Parameters').

D391N backbone signals were assigned following transposition of the WT assignment to the data. The 3D HNCO and HNCA data were sufficient to assign 82% of the backbone signals

(350/453) using the automated assignment software, CARA, and further peak centered in the software, Sparky.

***Cy1 and T1 (Holo and Loaded) Complexes.***

Cy1 and T1 interactions were evaluated using a differential isotopic labeling scheme for Cy1 and T1. Holo-T1 was expressed and purified using previously established protocols[12] with a $^{15}N/^{2}H/^{12}C$ or a $^{15}N/^{1}H/^{12}C$ labelling scheme, while Cy1 had either CDN or CDN-ILV labeling. For all complexes between T1 and Cy1, NMR titration buffer (50 mM ACES, 10 mM NaCl, 2 mM $MgCl_2$, pH 7.0) was used. Cy1 amide resonances were unperturbed in NMR titration buffer compared to the buffer used in signal assignments and structural calculation (20 mM sodium phosphate buffer, 10 mM NaCl, 1 mM EDTA, 5 mM DTT, pH 7, 0.05% w/v sodium azide). We established that addition of other buffer components salicylate, ATP, and $MgCl_2$ had no effect on the Cy1 or holo T1 spectra (data not shown).

The interaction between T1 (holo/loaded) and Cy1 was evaluated four times, with two instances in which T1 was loaded in the presence of Cy1 (*in situ*), while for the remaining two instances T1 was loaded externally and then added to Cy1. The initial *in situ* loading reaction used CDN labeled Cy1 (500 μM) and $^{15}N/^{1}H/^{12}C$ labeled holo T1 (560 μM) to interrogate binding. Holo-T1 was loaded *in situ* using 100 nM YbtE, 2 mM salicylate, and 2 mM ATP. In this, holo T1 was first complexed with Cy1, salicylate was then added, followed by ATP, and finally the loading reaction was initiated through the final addition of YbtE. After each addition of a reaction component, IDIS-TROSY spectra were collected to simultaneously evaluate both sample quality (T1 and Cy1) as well as to probe for any potential interactions between reagents and the two proteins. The loading of T1 through addition of YbtE was followed by IDIS-TROSY experiments.[45] We monitored the T1 holo to substrate-loaded conversion by observing the intensities of residue S52,[46] as this residue exhibits a new signal upon salicylate loading. This initial *in situ* reaction was used to obtain the holo T1 - Cy1 and loaded T1 - Cy1 HNCO spectra used to verify the absence of interactions between holo-T1 and Cy1, and to calculate the population of minor peaks in Cy1 in the presence of loaded T1, respectively. We then verified that we could reproduce the detection of minor peaks when T1 was loaded externally and added to Cy1. In this instance, CDN Cy1 (345 μM) and $^{15}N/^{1}H/^{12}C$ labeled loaded T1 (400 μM) were present in the final sample. Holo-T1 (at ~ 40 uM, 4 ml) was first loaded *in situ* by using 1 mM salicylate, 2 mM ATP, and 1 μM YbtE. The sample was then buffer exchanged into fresh NMR titration buffer using centrifugal filtration (3K molecular weight cut-off filter). Following buffer exchange, T1 was found loaded at 65 % (using the S52 signal); T1 and Cy1 were mixed to final concentrations of 400 μM and 345 μM, respectively. YbtE was present at 50 nM. An HNCO was acquired for loaded T1 in presence of Cy1, during which hydrolysis brought the loaded population to 51 %. The thioesterase SrfAD (500 nM), which enzymatically hydrolyzes the thioester bond of salicylate loaded to T1, was added to this sample to outcompete YbtE and a second HNCO was collected. This HNCO demonstrated that minor Cy1 signals seen in all experiments are only present when salicylate is loaded to T1 through a thioester bond and are not due to accumulation of chemical products (e.g. AMP, Sal-AMP). This spectrum, and that of free Cy1, were also used to monitor for signals that may indicate degradation. The SrfAD thioesterase expression and purification has been presented previously.[46]

We repeated the above experiments with minor variations, first to correlate the increase in intensities of Cy1 minor signals with T1 loading, and finally to help assign resonances of Cy1's minor conformer. We monitored an *in situ* loading reaction of U-$^{15}$N-$^{2}$H-$^{12}$C holo T1 (300 μM) in presence of CDN-ILV Cy1 (100 μM) and various concentrations of YbtE using time-shared HN-TROSY/HC-HSQCs acquired every 7 minutes over the course of 30 hours for a total of 190 spectra. This HN-TROSY/HC-HSQC is the 2D version of a previously published timeshared NOESY.[32] The signals of S52 and R77 in the HN-TROSY sub-spectrum of T1 are spectrally isolated from Cy1 signals and were used to monitor T1 loading. Cy1 minor signals were monitored for the most intense signals of Cy1 that gave minor signals, V237, W402, and T314. Concomitant HC-HSQC spectra provided Cy1 minor methyl signals for three spectrally isolated isoleucines I428, I60, and I236. T1 loading was started using 5 nM YbtE and stalled at 10% loading over the course of 7 hours. We then increased the YbtE concentration in the sample to 50 nM and time-shared HN-TROSY/HC-HSQC spectra were again acquired until approximately 80% of T1 was loaded. Finally, YbtE was increased to 200 nM and no additional increase in T1 loading was observed. Seven sets of 15 spectra were summed from that pool of spectra to increase the sensitivity and correlate loading of T1 with an increase in intensity of Cy1 minor peaks (Fig.3d-e and SI Fig. 5a). The population of the Cy1 minor conformer was calculated by

$$Population(Cy1_{minor}) = \frac{I_{minor}}{I_{major} + I_{minor}}$$

where I is the intensity of the major or minor peaks. The bars in Fig. 3 represent averages over the 6 Cy1 residues. The error reported is calculated through error propagation. We first calculated the error for the population of each residue, using the noise as the error for the intensities used in the equation above, and further used error propagation to calculate the error of the mean value.

Finally, we used a sample of CDN-ILV Cy1 (300 μM) presented to externally loaded T1 (395 μM) to acquire a 3D HNCO and 3D HNCA to assign Cy1 minor peaks. In preparation, 447 μM holo-T1 was loaded using 4 mM salicylate, 5 mM ATP, and 1 μM YbtE for one hour to achieve approximately 80% loading. This loaded T1 sample was then purified by size exclusion chromatography, concentrated using centrifugal filters, and mixed with CDN-ILV Cy1 to final concentrations of 395 μM and 300 μM for T1 and Cy1, respectively. T1 was observed loaded to 60% in this complex, and 20 nM YbtE was added to maintain similar amounts of loaded T1 throughout the two 3D acquisitions.

In addition to visual inspection of all NMR datasets, the integrity of the samples was monitored by SDS-PAGE of aliquots taken at different points of our experiments (see SI Fig. 6c).

***Assignment of Minor Cy1 Resonances.***

The signals of the minor conformer of Cy1 were assigned through multiple 3D datasets collected over the course of the reactions described above. Thus, $^{1}H$, $^{15}N$, $^{13}CO$, $^{13}C\alpha_i$, and $^{13}C\alpha_{i-1}$ chemical shifts could be assigned to minor peaks, except in 7 (out of 77) where sequential signals to $C^{\alpha}$ of the previous residue was missing. For most residues, the frequencies of $C^{\alpha}$ or CO or both resonances were similar for major and minor conformers. To account for possible degradation in Cy1, we inspected 3D HNCO spectra collected after unloading of T1 with the thioesterase SrfAD, as signals of the minor conformer would disappear following addition of SrfAD but degradation peaks would not. That is, signals of the minor state were assigned through a combination of spectroscopy (to assign them to residues) and biochemistry (to verify that they reported on the response of Cy1 towards loading of T1).

***In situ Loading of Holo-T1 in the Presence of Cy1 D391N.***

To monitor the allosteric response of Cy1 D391N with salicylate-loaded T1, we performed the *in situ* assay similar to that done with Cy1 WT. Briefly, 285 μM Cy1 D391N (CDN) was complexed with 386 μM holo-T1 ($^{15}$N) in titration buffer (50 mM ACES, 10 mM NaCl, 2 mM $MgCl_2$, 1 mM TCEP, pH 6.95 at room temperature) containing 1 mM salicylate and 2 mM ATP. 98 nM YbtE was added to start the loading reaction. Following confirmation of T1 loading with salicylate, 2D IDIS-TROSY and 3D HNCO data were collected on this complex over the course of 10 days. When needed, as monitored through 1D NMR spectra, ATP was spiked into the sample. Following HNCO data collection, 491 nM of the thioesterase SrfAD was added to unload T1 and another 3D HNCO dataset was collected.

For the loaded T1-Cy1 D391N complex, 3D HNCO and HNCA data were collected. After unloading of T1 by SrfAD, a 3D HNCO dataset was collected using the same parameters. All 3D HNCO and HNCA datasets were processed using NMRpipe with zero-filling of 512 points in the $^{15}$N and $^{13}$C dimensions using a cosine-squared bell function for apodization and linear prediction. For figures in which data that included HN-projections of 3D HNCO datasets, the 3D dataset was skyline projected using the Xeasy *project* script over the $^{13}$C dimension using points 58 to 505 such that $t_1$-noise from buffer components at the edges of the spectra did not obscure the projection. Negatively phased signals were contoured out in CARA to reduce the size of the files that would otherwise become too large for figure production due to $t_1$-noise from residual ATP signals. Further, 2D IDIS-TROSY spectra were collected between all 3D datasets to assess the quality of both T1 and Cy1 D391N and to ensure T1 attained a loaded state before each 3D dataset. Interleaved data was split in TopSpin 3.6.2 using the *split ipap* command.

To quantify the percentage of substrate-loading on holo-T1 in the complex, we computed the relative intensities of loaded T1 over the ratios of the holo and loaded T1 intensities as reported previously.[46] We averaged the population of loaded T1 over the following residues: S52, I53, R54, L59, and A77. The average population is reported in SI Fig. 9b, with the error as the standard deviation across these residues.

***Population Calculations and Determination of D391N Minor Peak Detection Limit.***

Cy1 WT minor populations were calculated as described in the Methods section**.** In Figure 4d, the Cy1 WT populations are denoted as a scatter plot with a box-and-whisker plot. In the box-and-whisker plot, the central line of the box denotes the median of all Cy1 WT populations with the $25^{th}$ and $75^{th}$ percentiles defining the edges of the box. The end of the whiskers denotes the extremes of the data after discarding outliers, that is, populations with more than 1.5 times the interquartile range. All figures were generated in MatLab 2020a.[47]

In spite of a comparable signal-to-noise in the D391N spectra, we could not assign four signals that we detected in D391N in presence of loaded-T1. These minor peaks subsequently disappeared upon addition of SrfAD, however, their assignment was still undetermined. As a result, we report the population of the Cy1 D391N minor peaks as a limit of detection. That is, we used the equation described in the methods section but replaced $I_{minor}$ with the noise for that position. We then took the average across all residues that displayed minor peaks in the spectra recorded with wild type Cy1. We report this limit of detection with the error as the standard deviation across all residues.

***Protein Dynamics.***

TROSY $^{15}N$ relaxation dispersion data[48,49] was acquired at 600 MHz and 950 MHz on a 600 μM sample of $^{1}$H-Me-δI-δ2Lγ2V-$^{2}$H-$^{15}$N Cy1. 13 points (950 MHz) and 15 points (600 MHz) including the reference point (for the $R_{2,eff}$ calculation) were acquired with frequencies ranging from 34.92 Hz to 961.54 Hz (950 MHz) and 29.07 – 1008.06 Hz (600 MHz). The analysis of relaxation dispersion profiles is described in detail in the Methods section of the main manuscript.

***Calculation of Chemical Shift Perturbations (CSPs):***

Backbone and Cy1 major/minor CSPs were calculated using $^{1}H^{N}$, $^{15}N$, and $^{13}CO$ chemical shifts from peak-centered 3D HNCO spectra. The magnitude of the CSPs (Δδ) were calculated using the following equation:[50]

$$\Delta\delta = \sqrt{\left[\frac{1}{3}\left\{\delta\left({}^{1}H^{N}\right)^{2} + \delta\left({}^{15}N\right)^{2} + \delta\left({}^{13}CO\right)^{2}\right\}\right]}$$

where δ is the chemical shift difference in ppm for a given nuclei (in parentheses). CSP calculations were performed using in-house Matlab scripts.

In Fig. 4a, we have mapped the D391N CSPs onto the medoid structure of Cy1. For the structure mapping, a color gradient was applied to emphasize CSPs above the median. In this, an in-house Python script was written to generate the color gradient utilizing the medoid structure in PyMOL 2.5.0. This gradient scale applies a grey to red color ramp, with grey being defined as a median CSP (or lower) and the brightest red as two standard deviations above the median. Here, the unnormalized median CSP was 0.035 and two standard deviations above the median was 0.248. Residues that were uniquely assigned in Cy1 D391N, but not in WT, were highlighted in orange to signify that they have a CSP, but whose magnitude is unknown. We note here that their assignment (although few in number) could be a result of more efficient back exchange in the D391N mutant or a considerable CSP from WT that positions the resonances in a region with overlap.

##### *Detection of Chemical Exchange using Hartmann-Hahn $R_{ex}$ Profiles.*

To compare the dynamics of Cy1 WT and D391N, we utilized the method of Wang et al., [37] which relies on estimating the exchange-contribution to relaxation of the individual components of $^{15}$N doublets. Here, a 600 μM NMR sample of Cy1 WT (CDN) and a 122 μM NMR sample of Cy1 D391N (CDN) in NMR buffer (20 mM sodium phosphate, 10 mM NaCl, 1 mM EDTA, and 5 mM DTT, pH 7 at room temperature) with 10% v/v $D_2O$ and 1% v/v DSS were used. Data was acquired at 600 MHz (see section 'NMR Experiments and Acquisition Parameters'). Following acquisition, the data was zero-filled to 1024 points in $^{15}$N, apodized using a cosine-squared bell function, and linear predicted. The final processed dataset was extracted over the amide region in the $^{1}$H dimension and subsequently analyzed in Sparky. The pulse sequences described in [37] were modified to include $^{13}$C decoupling via adiabatic Chirp inversion pulses in $^{15}$N encoding and long transverse relaxation periods to account for $^{13}$C labeling in the Cy1 samples. Water suppression utilizing a 3-9-19 block was used instead of excitation sculpting.

Analysis of the data for WT and D391N was conducted as described in [37] using in-house Matlab scripts. Briefly, the intensities of signals in the three different pulse sequences allow for estimation of the relaxation rates $R_2^{\alpha}$, $R_2^{\beta}$, and $R_1^{2HzNz}$, where $R_2^{\alpha}$ and $R_2^{\beta}$ are the transverse relaxation rates of the operators $H^{\alpha}N^{-}$ and $H^{\beta}N^{-}$, respectively, and $R_1^{2HzNz}$ is the relaxation rate of the longitudinal two-spin order $2H_zN_z$. The contribution to relaxation from exchange processes is obtained through:

$$R_{ex} = R_2^{\alpha} - (R_1^{2HzNz}/2) - \eta_{xy}(\kappa - 1)$$

in which $\eta_{xy}$, is the cross-correlation rate constant between the $^{15}N$ dipole-dipole and chemical shift anisotropy interactions, and the scaling factor κ is determined through the 5% trimmed mean of:

$$1 + \frac{(R_2^{\alpha} - R_1^{2HzNz}/2)}{\eta_{xy}}$$

Monte Carlo error analysis was performed over 300 steps using the noise as a variance, with the mean $R_{ex}$ and their standard deviations to the mean reported in Fig. 4.

***CAVER Tunnel Analysis of Cy1 NMR Ensemble.***

To assess heterogeneity of tunnels within each conformer of the Cy1 NMR ensemble, we used the CAVER 3.0.3[51] PyMol plug-in (Fig. 2a,d and SI Fig. 2c). As we sought to identify what would happen to tunnels in presence of structural fluctuations, we used a small probe radius so tunnels could be found even in presence of constrictions. For each of the 20 conformers in the ensemble, tunnel calculations were performed using a 0.7 Å probe radius, 4.0 Å shell radius, 4.0

Å shell depth, and a clustering threshold of 3.5. The center of the Cy1 tunnel cavity was used as a starting point, i.e. all residues were selected in each conformer and the cartesian coordinate center was calculated using the CAVER plug-in. As CAVER determines tunnels by starting at an optimized point and reaching the surface of the structure, in some Cy1 conformers, the probe radius was further decreased when putative donor and acceptor thiolation tunnel entrances were inaccessible at 0.7 Å. Specifically, State 12 and 18 used a 0.2 Å probe radius. In State 19, due to the limited accessibility at the acceptor T domain binding site, the starting point of the tunnel was set between residues 153 and 359 near the entrance formed by L20 and L16, and the parameters were changed to a 0.2 Å probe radius, a 6 Å shell depth, a 4 Å shell radius, and a clustering threshold of 3.5. In all tunnel calculations, several erroneous tunnels are determined throughout the structures due to the small probe radius, and we identified tunnels that align with those identified in open states, when the probe could be increased to 1.2 Å. With this protocol we could determine tunnels even when the donor (L20 and L16) and acceptor (α1 and α10) were closed to substrates.

***Thermal Denaturation Monitored by Circular Dichroism (CD) and Fluorescence.***

The apparent melting temperatures of WT and D391N Cy1 were assessed by thermal denaturation and monitored by CD on an AVIV Model 420 CD spectrometer. Cy1 WT and Cy1 D391N (unlabeled) were used at concentrations of 0.035 and 0.028 mg/mL, respectively, in NMR buffer (20 mM sodium phosphate, 10 mM NaCl, 1 mM EDTA, pH 7) with 5 μM TCEP and 0.0025% w/v $NaN_3$. Prior to thermal denaturation, CD spectra of each protein were recorded over a wavelength range from 195 to 260 nm (SI Fig. 8a) in intervals of 1.0 nm and averaging the signal over 1.0 seconds at 25.0 °C in a 1-cm quartz cell. Thermal denaturation was performed over a range of 20 to 69 °C with a temperature step of 1.00 °C, 60 seconds of equilibration time, 15 seconds of signal averaging, and monitoring the signal at 222.0 nm in a 1.0-cm pathlength quartz cuvette. To ensure sample homogeneity, each sample was stirred during the measurements. After collection of at least 10 points in the unfolded baseline, the samples were cooled to 25.0 °C at a rate of 5 °C $min^{-1}$ and a spectrum of each sample was collected again to assess reversibility.

Both WT and D391N samples exhibited a two-state unfolding transition. The raw CD data was corrected for concentration dependence by converting to units of molar ellipticity using an extinction coefficient of 88,265 $M^{-1}$ $cm^{-1}$ and 453 residues in Cy1. Extraction of apparent melting temperatures (SI Fig. 8c) for each sample was determined by fitting the data to a two-state model using the following equation:

$$\alpha_{obs} = \alpha_N + (\alpha_D - \alpha_N)\left(\frac{e^{\Delta \mathrm{H(T-Tm)/RTTm}}}{e^{\Delta \mathrm{H(T-Tm)/RTTm}} + 1}\right)$$

where $\alpha_{obs}$ is the CD signal at 222.0 nm, T is temperature (Kelvin), $\alpha_N$ and $\alpha_D$ are linear fits to the native (folded) and denatured baselines respectively, ΔH is the apparent enthalpy (kJ $mol^{-1}$), $T_m$ is the apparent melting temperature (Kelvin), and R is the universal gas constant (0.0083145 kJ $K^{-1}$ $mol^{-1}$). The parameters were fit using a non-linear model in Wolfram Mathematica 12.0. Reported errors for fitted parameters were generated by computing the standard error of the fit.

Although the apparent enthalpy was parameterized independent of temperature, it is critical to note that the thermal unfolding of both samples was irreversible.

##### *Measurement of $^1H,^{15}N$ Residual Dipolar Couplings (RDCs).*

The $^1H,^{15}N$ RDC values for wild type Cy1 were measured on a $^2H,^{13}C,^{15}N$-Cy1-H6 sample aligned using 12 mg/mL of Pf1 phage (ASLA Biotech) and a deuterium splitting of ~11.4 Hz. RDCs were calculated as the difference between J+D measured on the aligned sample and J value measured on the isotropic sample. The J+D and J values are obtained as twice the difference between the nitrogen chemical shifts (measured in Hz) in non-TROSY and TROSY versions of the 3D-HNCO. Trosy and non-Trosy versions of 3D-HNCO spectra were acquired in an interleaved manner using NUS protocols (65 points in $^{15}N$, 35 points in $^{13}C$, and 30 % of sampling giving rise to 832 complex points) (SI Table S4). 3D spectra were processed with zero-filling to 512 points in the $^{15}N$ dimension and 256 points in the $^{13}C$ dimension. The spectra were initially loaded in CARA and the peaks were centred manually. Using in-house scripts, the peak lists were exported to tab (NMRPipe) format and were used as input to run NMRPipe peak fitting program nlinLS. The chemical shift values of the peaks were allowed to vary during the nlinLS fits to finetune the centering of the peaks and obtain an estimate of the error in chemical shift. nlinLS returned peak positions and linewidths that were fit and associated errors in the units of points for each spectrum. The errors were converted into Hz and were propagated during the calculation of RDC values. Simulated 3D-HNCO spectra were created using the NMRPipe script “simSpecND” using the chemical shift and linewidth values returned by nlinLS. The simulated 3D-HNCO spectra were superposed on the acquired spectra in CcpNMR analysis and peak overlays were scanned through to visually check the goodness of the fit. A few iterations were carried out by changing the initial input parameters to improve the fits. A total of 333 RDCs were measured at this initial stage. RDCs involving $^{15}N$ nuclei with order parameter $S^2 <$

0.7 (determined using TALOS+[52]) and those exhibiting chemical exchange in the RD-CPMG experiments were excluded from the restraint list resulting in 258 RDCs. The RDC of a residue with $S^2$ of 0.692 was rescued and included in calculations.

***Structure restraints and structure calculation.***

Distance restraints were obtained from unambiguously assigned inter-proton correlations in a set of $^{13}C$ and $^{15}N$ edited 3D-NOESY spectra (SI table S1). 4D-NOESY spectra were used to resolve ambiguities. Time-shared acquisition of $^{13}C$ and $^{15}N$ edited NOESY spectra with NOESY in the acquired dimension[32] proved critical for obtaining optimal resolution in the NOESY dimension for unambiguous assignment of distance restraints, particularly at 950 MHz where reaching such a resolution in the indirect dimension would require prohibitive experimental times. Dihedral angle restraints were prepared using a consensus between TALOS+[52] and CSI 3.0.[53] Dihedral angle restraints for 7 residues in α4 and 5 residues in β11, with either incomplete chemical shift assignments or weak signals that could not provide constraints, were prepared based on a preliminary RASREC CS-Rosetta structure model.[54] Secondary structure elements were identified based on NOE patterns and chemical shift index from CSI 3.0, and hydrogen bond restraints were included for related residues. Hydrogen bond restraints for 7 residues in α4 and 5 residues in β11 were also based on the RASREC CS-Rosetta structure model which used NMR chemical shifts, NOEs and RDCs alongside several optimization strategies to compute near-native structures. The sampling of accurate folds was carried out across multiple stages. The early stages focussed on exploration of beta-sheet topologies followed by sampling of fragments from high-resolution X-ray structures and low-resolution models to obtain final folds which underwent refinement to produce high-resolution models.

Structure calculations were carried out using CYANA 3.98.[55-57] An initial structure bundle of Cy1 was calculated using accurately calibrated distance restraints obtained from a pair of $^{13}C$ and $^{15}N$ edited time-shared 3D-NOESY spectra (mixing time = 40 ms) recorded at 950 MHz on a Cy1 sample with stereospecific labelling of methyl groups of Leu and Val. The short mixing time and stereo-specific methyl labels ensured accurate volume integration with minimal contamination from spin diffusion. A total of 500 structures were calculated from 1187 distance constraints (and including 810 dihedral angle restraints and 293 hydrogen bond restraints), and the 50 structures with lowest energy were chosen for further analysis. This initial structure bundle had a backbone RMSD of 1.96 Å for structured regions. To identify a representative conformation of Cy1, the structures in the bundle were clustered based on pair-wise RMSD distance matrix, and the model closest to the mean of the largest cluster was selected as most representative of the conformation captured by NMR. An additional 1,002 distance restraints were identified from all NOESY spectra with longer mixing times. These restraints were selected from the complete set of experimental data (excluding those at 40 ms) and binned into groups with upper limits at 3, 5, 7, and 9 Å such that they were in agreement with the initial model obtained with distances calibrated at 40 ms mixing time. NOESY peak assignments were manually curated and refined iteratively over several CYANA runs to remove violations due to overlap or erroneous assignments. 173 RDC restraints could be included without violations in the final stages of structure calculation in addition to all 2189 distance restraints, 810 dihedral angle restraints, and 293 hydrogen bond restraints. The 50 structures with lowest energy from a total of 500 structures were selected, refined in explicit solvent using CNS,[58] and models with short contacts or improper secondary structure elements were screened out. The 20 models with lowest energy were selected to make the final bundle, with no distance violations larger than 0.5Å present and a pairwise root-mean-square standard deviation of 1.5 Å for ordered residues and 1.2 Å for those in secondary-structured regions.

***Consensus sequences, threaded conformational models, and electrostatic potential map calculations.***

Consensus sequences were obtained from Rausch et al.[59] who conducted a phylogenetic study on C domain functional subtypes. For each sub-type, the consensus sequences were extracted from the sequence logo files provided in [59], narrow width stacks (position with many gaps) were screened out. Only the tallest symbol indicating the most conserved residue at that position were used in the consensus sequences. All the resulting sequences were then aligned using a combination of structural (using PDBs) and sequence alignment (consensus sequences from $^{L}C_{L}$, cyclization, C starter, $^{D}C_{L}$, and dual epimerization/condensation domains) using PROMALS3D[60]. The resulting multiple sequence alignment along with choice of chosen C-domain sequence and conformation (pdb ids: 4JN3, 6P1J, 1L5A, and 5T3D) were used to generate conformational models in SI Fig. 10d, using the SWISS-MODEL[61] target-template alignment. For example, the conformation in SI Fig. 10d, row 1, column 1 was generated using the sequence alignment between the $^{L}C_{L}$ consensus sequence and that of pdb id 4JN3, such that the resulting conformation has the $^{L}C_{L}$ consensus sequence threaded onto the conformation of 4JN3. For model comparison in SI Fig.10 b and c, the C-term sub-domains were aligned; the regions corresponding to loops 16, 20, and β11 were left out of this alignment. The electrostatic potential maps were then calculated using these structural models using the Adaptive Poisson

Boltzmann Solver[62] as part of Pymol plugins. The solvent excluded surface (Connolly surface) was used for the calculation.

12 Goodrich, A. C., Harden, B. J. & Frueh, D. P. Solution Structure of a Nonribosomal Peptide Synthetase Carrier Protein Loaded with Its Substrate Reveals Transient, Well-

Defined Contacts. *Journal of the American Chemical Society* **137**, 12100-12109, doi:10.1021/jacs.5b07772 (2015).
13 Keating, T. A., Marshall, C. G., Walsh, C. T. & Keating, A. E. The structure of VibH represents nonribosomal peptide synthetase condensation, cyclization and epimerization domains. *Nature structural biology* **9**, 522-526, doi:10.1038/nsb810 (2002).
14 Tan, K. *et al.* Structures of teixobactin-producing nonribosomal peptide synthetase condensation and adenylation domains. *Current Research in Structural Biology* **2**, 14-24, doi:https://doi.org/10.1016/j.crstbi.2020.01.002 (2020).
15 Drake, E. J. *et al.* Structures of two distinct conformations of holo-non-ribosomal peptide synthetases. *Nature* **529**, 235-238, doi:10.1038/nature16163 (2016).
16 Tanovic, A., Samel, S. A., Essen, L. O. & Marahiel, M. A. Crystal structure of the termination module of a nonribosomal peptide synthetase. *Science* **321**, 659-663 (2008).
17 Frueh, D. P. *et al.* Dynamic thiolation-thioesterase structure of a non-ribosomal peptide synthetase. *Nature* **454**, 903-906, doi:10.1038/nature07162 (2008).
18 Miller, B. R., Drake, E. J., Shi, C., Aldrich, C. C. & Gulick, A. M. Structures of a Nonribosomal Peptide Synthetase Module Bound to MbtH-like Proteins Support a Highly Dynamic Domain Architecture. *J Biol Chem* **291**, 22559-22571, doi:10.1074/jbc.M116.746297 (2016).
19 Reimer, J. M., Aloise, M. N., Harrison, P. M. & Martin Schmeing, T. Synthetic cycle of the initiation module of a formylating nonribosomal peptide synthetase. *Nature* **529**, 239-242, doi:10.1038/nature16503 http://www.nature.com/nature/journal/v529/n7585/abs/nature16503.html#supplementary-information (2016).
20 Reimer, J. M. *et al.* Structural insight into a novel formyltransferase and evolution to a nonribosomal peptide synthetase tailoring domain. *ACS Chemical Biology*, doi:10.1021/acschembio.8b00739 (2018).
21 Tarry, M. J., Haque, A. S., Bui, K. H. & Schmeing, T. M. X-Ray Crystallography and Electron Microscopy of Cross- and Multi-Module Nonribosomal Peptide Synthetase Proteins Reveal a Flexible Architecture. *Structure*, doi:https://doi.org/10.1016/j.str.2017.03.014 (2017).
22 Tufar, P. *et al.* Crystal Structure of a PCP/Sfp Complex Reveals the Structural Basis for Carrier Protein Posttranslational Modification. *Chem. Biol.* **21**, 552-562, doi:DOI 10.1016/j.chembiol.2014.02.014 (2014).
23 Goodrich, A. C., Meyers, D. J. & Frueh, D. P. Molecular impact of covalent modifications on nonribosomal peptide synthetase carrier protein communication. *J Biol Chem* **292**, 10002-10013, doi:10.1074/jbc.M116.766220 (2017).
24 Qiao, C., Wilson, D. J., Bennett, E. M. & Aldrich, C. C. A mechanism-based aryl carrier protein/thiolation domain affinity probe. *J. Am. Chem. Soc.* **129**, 6350-6351, doi:10.1021/ja069201e (2007).
25 Sundlov, Jesse A., Shi, C., Wilson, Daniel J., Aldrich, Courtney C. & Gulick, Andrew M. Structural and Functional Investigation of the Intermolecular Interaction between NRPS Adenylation and Carrier Protein Domains. *Chem. Biol.* **19**, 188-198, doi:10.1016/j.chembiol.2011.11.013 (2012).
26 Shi, C., Miller, B. R., Alexander, E. M., Gulick, A. M. & Aldrich, C. C. Design, Synthesis, and Biophysical Evaluation of Mechanism-Based Probes for Condensation

Domains of Nonribosomal Peptide Synthetases. *ACS Chem Biol* **15**, 1813-1819, doi:10.1021/acschembio.0c00411 (2020).
27 Bloudoff, K. & Schmeing, T. M. Structural and functional aspects of the nonribosomal peptide synthetase condensation domain superfamily: discovery, dissection and diversity. *Bba-Proteins Proteom* **1865**, 1587-1604, doi:10.1016/j.bbapap.2017.05.010 (2017).
28 Schoppet, M. *et al.* The biosynthetic implications of late-stage condensation domain selectivity during glycopeptide antibiotic biosynthesis. *Chemical science* **10**, 118-133, doi:10.1039/c8sc03530j (2019).
29 Kosol, S. *et al.* Structural basis for chain release from the enacyloxin polyketide synthase. *Nature chemistry* **11**, 913-923, doi:10.1038/s41557-019-0335-5 (2019).
30 Samel, S. A., Schoenafinger, G., Knappe, T. A., Marahiel, M. A. & Essen, L. O. Structural and functional insights into a Peptide bond-forming bidomain from a nonribosomal Peptide synthetase. *Structure* **15**, 781-792 (2007).
31 Mishra, S. H. & Frueh, D. P. Assignment of methyl NMR resonances of a 52 kDa protein with residue-specific 4D correlation maps. *J Biomol NMR* **62**, 281-290, doi:10.1007/s10858-015-9943-6 (2015).
32 Mishra, S. H., Harden, B. J. & Frueh, D. P. A 3D time-shared NOESY experiment designed to provide optimal resolution for accurate assignment of NMR distance restraints in large proteins. *J Biomol NMR* **60**, 265-274, doi:10.1007/s10858-014-9873-8 (2014).
33 Harden, B. J., Mishra, S. H. & Frueh, D. P. Effortless assignment with 4D covariance sequential correlation maps. *J. Magn. Reson.* **260**, 83-88, doi:10.1016/j.jmr.2015.09.007 (2015).
34 Kancherla, A. K., Marincin, K. A., Mishra, S. H. & Frueh, D. P. Minimizing Pervasive Artifacts in 4D Covariance Maps for Protein Side Chain NMR Assignments. *J Phys Chem A* **125**, 8313-8323, doi:10.1021/acs.jpca.1c05507 (2021).
35 Goto, N. K., Gardner, K. H., Mueller, G. A., Willis, R. C. & Kay, L. E. A robust and cost-effective method for the production of Val, Leu, Ile (delta 1) methyl-protonated 15N-, 13C-, 2H-labeled proteins. *J. Biomol. NMR* **13**, 369-374 (1999).
36 Liu, H. & Naismith, J. H. An efficient one-step site-directed deletion, insertion, single and multiple-site plasmid mutagenesis protocol. *BMC Biotechnol.* **8**, 91, doi:10.1186/1472-6750-8-91 (2008).
37 Wang, C., Rance, M. & Palmer, A. G., 3rd. Mapping chemical exchange in proteins with MW > 50 kD. *Journal of the American Chemical Society* **125**, 8968-8969, doi:10.1021/ja035139z (2003).
38 Delaglio, F. *et al.* NMRPipe a Multidimensional Spectra Processing System Based on UNIX Pipes. *JBNMR* **6**, 277-293 (1995).
39 Keller, R. L. J. *The Computer Aided Resonance Assignment Tutorial*. (Cantina Verlag, 2004).
40 Kneller, T. D. G. a. D. G. SPARKY 3. *University of California, San Francisco*.
41 Kancherla, A. K. & Frueh, D. P. Covariance nuclear magnetic resonance methods for obtaining protein assignments and novel correlations. *Concepts Magn Reson Part A Bridg Educ Res* **46A**, doi:10.1002/cmr.a.21437 (2017).

42 Harden, B. J. & Frueh, D. P. Covariance NMR Processing and Analysis for Protein Assignment. *Methods Mol Biol* **1688**, 353-373, doi:10.1007/978-1-4939-7386-6_16 (2018).

43 Hyberts, S. G., Milbradt, A. G., Wagner, A. B., Arthanari, H. & Wagner, G. Application of iterative soft thresholding for fast reconstruction of NMR data non-uniformly sampled with multidimensional Poisson Gap scheduling. *J. Biomol. NMR* **52**, 315-327, doi:10.1007/s10858-012-9611-z (2012).

44 Vallurupalli, P., Bouvignies, G. & Kay, L. E. Studying "invisible" excited protein states in slow exchange with a major state conformation. *Journal of the American Chemical Society* **134**, 8148-8161, doi:10.1021/ja3001419 (2012).

45 Golovanov, A. P., Blankley, R. T., Avis, J. M. & Bermel, W. Isotopically discriminated NMR spectroscopy: a tool for investigating complex protein interactions in vitro. *J Am Chem Soc* **129**, 6528-6535 (2007).

46 Goodrich, A. C. & Frueh, D. P. A nuclear magnetic resonance method for probing molecular influences of substrate loading in nonribosomal peptide synthetase carrier proteins. *Biochemistry* **54**, 1154-1156, doi:10.1021/bi501433r (2015).

47 Matlab. *9.7.0.1190202 (R2019b)*. (Math Works Inc., 2018).

48 Loria, J. P., Rance, M. & Palmer, A. G., 3rd. A TROSY CPMG sequence for characterizing chemical exchange in large proteins. *J Biomol NMR* **15**, 151-155, doi:10.1023/a:1008355631073 (1999).

49 Hansen, D. F., Vallurupalli, P. & Kay, L. E. An Improved 15N Relaxation Dispersion Experiment for the Measurement of Millisecond Time-Scale Dynamics in Proteins†. *The Journal of Physical Chemistry B* **112**, 5898-5904, doi:10.1021/jp074793o (2008).

50 Terada, T. *et al.* Nuclear magnetic resonance and molecular dynamics studies on the interactions of the Ras-binding domain of Raf-1 with wild-type and mutant Ras proteins. *J Mol Biol* **286**, 219-232, doi:10.1006/jmbi.1998.2472 (1999).

51 Chovancova, E. *et al.* CAVER 3.0: a tool for the analysis of transport pathways in dynamic protein structures. *PLoS Comput Biol* **8**, e1002708, doi:10.1371/journal.pcbi.1002708 (2012).

52 Shen, Y., Delaglio, F., Cornilescu, G. & Bax, A. TALOS+: a hybrid method for predicting protein backbone torsion angles from NMR chemical shifts. *J Biomol NMR* **44**, 213-223, doi:10.1007/s10858-009-9333-z (2009).

53 Hafsa, N. E., Arndt, D. & Wishart, D. S. CSI 3.0: a web server for identifying secondary and super-secondary structure in proteins using NMR chemical shifts. *Nucleic Acids Res* **43**, W370-377, doi:10.1093/nar/gkv494 (2015).

54 Lange, O. F. *et al.* Determination of solution structures of proteins up to 40 kDa using CS-Rosetta with sparse NMR data from deuterated samples. *Proc Natl Acad Sci U S A* **109**, 10873-10878, doi:10.1073/pnas.1203013109 (2012).

55 Guntert, P., Mumenthaler, C. & Wüthrich, K. Torsion angle dynamics for NMR structure calculation with the new program DYANA. *J Mol Biol* **273**, 283-298 (1997).

56 Guntert, P. Automated NMR structure calculation with CYANA. *Methods Mol Biol* **278**, 353-378 (2004).

57 Schmidt, E. & Guntert, P. Automated structure determination from NMR spectra. *Methods Mol Biol* **1261**, 303-329, doi:10.1007/978-1-4939-2230-7_16 (2015).

58 Brunger, A. T. Version 1.2 of the Crystallography and NMR system. *Nat Protoc* **2**, 2728-2733, doi:10.1038/nprot.2007.406 (2007).
59 Rausch, C., Hoof, I., Weber, T., Wohlleben, W. & Huson, D. H. Phylogenetic analysis of condensation domains in NRPS sheds light on their functional evolution. *BMC Evol. Biol.* **7**, 78, doi:10.1186/1471-2148-7-78 (2007).
60 Pei, J. M. & Grishin, N. V. PROMALS3D: Multiple Protein Sequence Alignment Enhanced with Evolutionary and Three-Dimensional Structural Information. *Multiple Sequence Alignment Methods* **1079**, 263-271, doi:10.1007/978-1-62703-646-7_17 (2014).
61 Bienert, S. *et al.* The SWISS-MODEL Repository-new features and functionality. *Nucleic Acids Res.* **45**, D313-D319, doi:10.1093/nar/gkw1132 (2017).
62 Jurrus, E. *et al.* Improvements to the APBS biomolecular solvation software suite. *Protein Sci* **27**, 112-128, doi:10.1002/pro.3280 (2018).
